# LIMIX: genetic analysis of multiple traits

**DOI:** 10.1101/003905

**Authors:** Christoph Lippert, Franceso Paolo Casale, Barbara Rakitsch, Oliver Stegle

## Abstract

Multi-trait mixed models have emerged as a promising approach for joint analyses of multiple traits. In principle, the mixed model framework is remarkably general. However, current methods implement only a very specific range of tasks to optimize the necessary computations. Here, we present a multi-trait modeling framework that is versatile and fast: LIMIX enables to flexibly adapt mixed models for a broad range of applications with different observed and hidden covariates, and variable study designs. To highlight the novel modeling aspects of LIMIX we performed three vastly different genetic studies: joint GWAS of correlated blood lipid phenotypes, joint analysis of the expression levels of the multiple transcript-isoforms of a gene, and pathway-based modeling of molecular traits across environments. In these applications we show that LIMIX increases GWAS power and phenotype prediction accuracy, in particular when integrating stepwise multi-locus regression into multi-trait models, and when analyzing large numbers of traits. An open source implementation of LIMIX is freely available at: https://github.com/PMBio/limix.

## 1 Introduction

The goal of genetic analyses of quantitative traits is to determine the sources of phenotypic variation, attributing it to effects of single loci, polygenic background and confounding factors. Linear mixed models (LMMs) have emerged as a powerful approach for this purpose and have been successfully applied to genome-wide association studies (GWAS) in structured populations [1, 2, 3, 4], quantification of the proportion of heritability that can be attributed to sets of SNPs [5, 6], prediction of phenotype from genotype [7], and to account for hidden confounding factors, for example in expression quantitative trait loci (eQTL) mapping [8, 9]. Recently, there has been increasing interest in methods that combine multiple traits that are correlated due to shared (but hidden) genetic or non-genetic influences in a single model. By properly modeling this trait correlation, a study can yield increased power to detect genuine genetic associations and better interpretation. Consequently, multi-trait LMMs that account for such correlations have been proposed, first in the field of animal breeding [10] and, more recently, also in the context of GWAS [11, 12]. In genetics, the use of general-purpose mixed modeling tools is rare (e.g. ref. [13]), mainly because these lack the computational efficiency that is required to handle larger sample sizes. Instead, a number of rigid and highly specialized tools have been developed that speed up the specific computations that are needed for a particular task, such as GWAS testing [1, 2, 3, 4, 12], or variance component estimation [5, 6].

However, because different multi-trait studies tend to have variable study designs, methods have to be flexible enough to adapt the model, but still remain tractable for larger datasets. Multiple trait-variates can for example correspond to repeated measurements of a single phenotype over time [14], in different tissues [15, 16], or under different environments [17, 18], but also to different phenotypes such as expression values for multiple genes [17, 19, 20, 18], or the variates of a blood profile [21]. Moreover, practical analyses often suggest iterative changes to the model, both to increase power and to test for specific genetic architectures. Starting from simple models of single traits, these can be extended to multi-trait models and subsequently to stepwise multiple loci models [22]. At each stage, it is important to adjust model parameters, and to objectively guide analysis decisions by rigorously comparing alternative models. This will require model checking strategies, methods for robust parameter estimation, as well as to address statistical problems such as overfitting.

Here we present LIMIX, a multi-trait mixed modeling framework that has the flexibility to perform a broad range of genetic analyses in one tool, without sacrificing computational efficiency. LIMIX enables modeling genetic or environmental factors by combining different fixed effects, sample covariances and trait covariances. Thanks to sophisticated parameter regularization LIMIX can robustly infer complex covariances between large numbers of traits without overfitting, thereby enabling joint analysis of tens or even hundreds of phenotypes, substantially more than previous methods. Finally, LIMIX is computationally efficient, utilizing speedups to circumvent unnecessary computations in multi-trait modeling that scale as the cube of the number of traits [23, 24, 12]. At the same time, LIMIX extends efficient association tests [3, 4] to multi-trait association tests for flexible genetic designs. We analyzed three very different datasets to highlight novel modeling aspects of LIMIX. We show how to combine multi-trait modeling and stepwise multi-locus association analyses by considering correlated blood lipid traits from the Northern Finland Birth Cohort 1996 (NFBC1966) [21]. Next, we jointly analyze abundance estimates of multiple transcript-isoforms of the same gene, estimated from RNA-Seq in human [20], thereby improving detection power and interpretation of the genetic architecture of isoform expression. We also demonstrate how multi-trait LMMs can readily be extended to the joint genetic analysis of entire pathways consisting of dozens of genes, both within and between environments [17]. And finally, we assess alternative models in terms of their ability to predict phenotype from genotype.

## 2 Results

### Improved flexibility and efficiency in multi-trait LMMs

Similar to genetic studies of single phenotypes using established univariate LMMs, LIMIX represents each phenotype as the sum of a number of fixed effects and random effects. Depending on the application, LIMIX allows free specification of each fixed and random effect, both for univariate and multi-trait models. For example, in multi-trait GWAS covariates and the single nucleotide polymorphism (SNP) tested are modeled as fixed effects and background variation, such as population structure and noise, as random effects [3, 4, 22, 12] that are modulated using an additional trait-trait covariance matrix.

Unlike previous computationally efficient single-trait and multi-trait LMMs [3, 4, 12, 11], models defined in LIMIX are not restricted to rigid designs to achieve speedups. Instead, models can be built freely and LIMIX automatically determines possible computational speedups [3, 4, 23, 24, 12] depending on the model specification and analysis task.

### Combining multi-trait with multi-locus models improves GWAS of lipid related traits

Four lipid-related traits in the NFBC1966 [11, 22, 12] have been analyzed using joint association tests of single loci to multiple-traits [11, 12] and in a separate study using models involving multiple loci effects on single traits [22]. Here, we revisit these data to show the advantages of combining these two models into a single multiple-trait and multiple-loci analysis (see Online Methods). This combination enables to explicitly test for allelic heterogeneity [22], i.e. multiple independent genetic signals from loci that are in linkage disequilibrium (LD), across different traits. Additionally, we show how additional tests for alternative genetic designs enable further interpretation of multiple-trait associations.

A single-locus association scan using a four-degrees-of-freedom any effect test across all phenotypes, similar to [12], yielded significant associations at ten different loci (*α* = 5 × 10^−8^, λ_*GC*_ = 0.99, Supplementary Table 1); see top panel in Figure 1a. Next, we compared these associations to the joint multi-trait multi-locus model. At an inclusion cutoff of 5 × 10^−8^ the multi-locus model uncovered 14 independent genetic effects, uncovering four independent secondary effects at four of the ten associated loci. Multiple SNPs selected by the model for each of the CRP, LIPC and CETP genes and the CEACAM16-TOMM40-APOE region suggest the presence of genetic heterogeneity in these well-characterized regions, which is in line with previous findings [25]. Moreover, at two of the regions, CRP and CEACAM16-TOMM40-APOE, the secondary SNP selected by the model was not significant in the single-locus association test, indicating an increase in GWAS power (An example is shown in Figure 1b lower panel, Supplementary Table 1).

**Figure 1.**
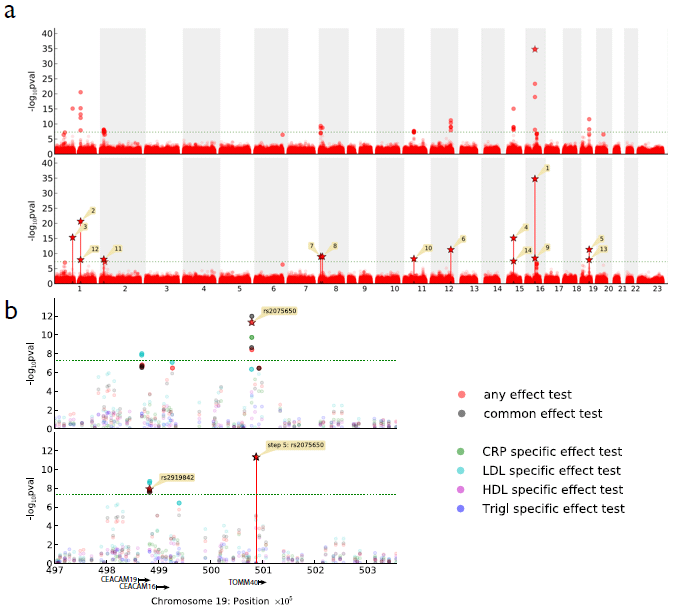
Multi-trait GWAS on four correlated phenotypes from the Northern Finish Birth Cohort (NFBC1966) [21]. **a**, upper panel: Manhattan plot of *P*-values from an any effects test in a multi-trait LMM. 34 SNPs have significant associations at 10 distinct loci (*α* = 5 × 10^−8^, *λ*_*G*_*C* = 0.99). lower panel: Manhattan plot showing the results of the novel multi-trait multi-locus LMM, using an any effects test. 14 SNPs with distinct genetic signals are included, each marked with a vertical bar. The numbered stars mark the *P*-values at the respective iteration of inclusion. **b**, Example zoom-in for a region with two independent effects included in the multi-locus LMM. Shown are several different association tests for the CEACAM-TOMM40 region before (upper panel) and after (lower panel) inclusion of the lead SNP (step 5, rs2075650) into the multi-trait multi-locus LMM. While SNPs that are associated purely due to LD to the primary associated SNP become insignificant after inclusion, a secondary association locus around rs2919842 (step 13) becomes more significant, pushing it above the genome-wide significance threshold for the any effect test (red) and the common effect test (black) (see also Supplementary Table 1).

To further tease apart effects that are shared between traits from trait-specific effects, we complemented the analysis with one specific effect test per phenotype and a common effect test across all phenotypes (see Online Methods). All tests were well-calibrated (see Supplementary Table 1 for λ_*GC*_ and *Q* – *Q*-plots in Supplementary Figure 1). The common effect test identified significant associations at 9 of the 10 loci detected by the any effect multi-locus test—all but the LIPC gene and a region on chromosome 8 between PPP1R3B and TNKS that had been associated with LDL levels in a larger study [26]. Notably, the phenotype-specific effect tests yielded significant associations at six out of the common effect loci, suggesting that many effects have both a common and specific component. This trend is also visible in independent single-trait analyses (Supplementary Table 1), suggesting that pleiotropic effects across all four phenotypes are scarce, unlike pairwise pleiotropy previously reported on the same dataset [11].

### Statistical tests for dissecting isoform-specific genetic regulation of gene expression

Association analyses for alternative genetic designs in LIMIX can also be applied to tease apart common from differential expression regulation of multiple isoform variants of single genes. Here, we analyzed deep RNA-Seq profiles for 464 HapMap samples [20], where we estimated the abundance of multiple isoforms for individual genes [27] (see Online Methods). We considered 9,246 genes with at least two expressed isoforms and used alternative LMMs to test for *cis* associations between proximal sequence variants (± 1mb around genes) and alternative isoform levels of these genes. Hidden confounding factors (beyond population structure) are a major concern in eQTL analyses [28, 29]. Consequently, we used LIMIX to estimate an extended sample-to-sample covariance that captures hidden confounding (similar to the PANAMA model [9]; see Supplementary Figure 2). As expected from previous results for single trait LMMs [9], accounting for hidden confounding increased power in multiple trait LMMs. Moreover, the observed power increase was even marginally larger in the multiple trait case than in the single trait case (Figure 2a; see Supplementary Table 2 for a list of significant associations).

**Figure 2.**
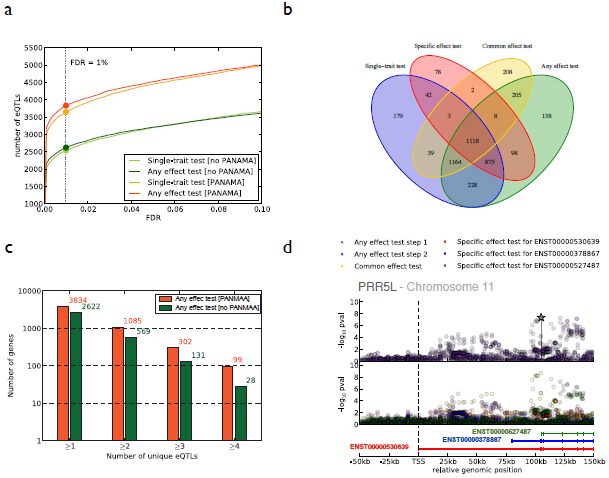
Isoform-specific eQTL analysis in the human eQTL dataset [20], considering 9,246 genes with at least two isoforms. **a**, Comparison of the number of genes with at least one significant *cis* eQTL (1mb upstream or downstream of each gene) as a function of the permitted false discovery rate (FDR) and for different association tests and models. Considered were an independent association test on each isoform (single-trait test) and a multi-trait LMM using the any effect test, both with and without accounting for hidden confounding factors (PANAMA). **b**, Overlap of genes having at least one significant *cis* association at an FDR cutoff of 1% when considering alternative tests in a multi-trait LMM: any effect tests, common effect tests and (isoform-)specific effect tests. **c**, Number of independent *cis* eQTLs (at 1% FDR), identified using the multi-locus multi-trait model, either with or without confounding correction (PANAMA). Bar plots denote the cumulative number of genes with at least (1, 2, 3 or 4) distinct *cis* eQTLs. **d**, Manhattan plot for a selected region around the gene PRR5L. The multi-trait multi-locus LMM applied to the region identified two independent associations. upper panel: *P*-values from an any effect test before inclusion of the lead SNP are shown in grey. *P*-values from an any effect test after inclusion of the lead SNP (grey asterisk) are shown in pink. lower panel: *P*-values from a common effect test and (isoform-)specific effect tests indicate that the lead association has a gene-level effect whereas the secondary association is specific to one isoform (ENST00000527487).

At a nominal significance level of FDR ≤ 1% (using *Q*-values [30]), the multi-trait model identified 3,648 genes (39.5%) with at least one eQTL in *cis*, 4.8% more than a single-trait LMM applied to each isoform independently (3,834 genes with an eQTL in *cis*, Figure 2a). Additionally, multi-trait testing enabled classifying individual eQTLs into genetic effects that affect whole-gene expression (common effect test eQTLs) and genetic effects that are isoform-specific (specific effect test eQTLs) (Figure 2b, Supplementary Figure 3).

Notably, for 29.4% of the genes with a *cis* eQTL (relative to any effect test, Figure 2b) the multi-trait model identified both an isoform-specific eQTL and a common effect eQTL in the same region, suggesting the presence of multiple independent genetic effects within single genes. To investigate this, we performed multi-locus analysis in the *cis* region of each gene jointly across all isoforms, yielding two or more independent associations (test statistics were calibrated; Supplementary Figure 4) for almost a third of all genes with an eQTL (28.3 %) and three or more independent associations for 27.8% of these (Figure 2c). A good example is PRR5L, a gene with four expressed isoforms (out of twenty-one annotated ones), where two unique eQTLs could be detected. The first lead eQTL corresponds to a common effect and the secondary eQTL was specific to one of the isoforms (Figure 2d).

Genome-wide, both the primary eQTLs and the independent higher-order associations all tended to occur near the transcription start site, which gives confidence that the multi-locus associations found are genuine [28, 20] (Supplementary Figure 5).

### Variance component models to characterize genotype-environment interactions on gene expression

Similar to the definition of flexible testing designs for fixed effects, LIMIX enables constructing interpretable designs for random effects. To show this, we revisited a yeast genetics dataset where gene expression levels have been measured under two conditions [17]. We demonstrate how to partition phenotypic variance, not only across genetic regions [6], but also within and between environments.

For this application, we considered an LMM that estimates components of expression variability for environmental effects, *cis* genetic effects, *trans* genetic effects, and their respective interactions (*cis* GxE, *trans* GxE) (see Online Methods). Unlike related methods for cross-tissue heritability estimation [15], we tease apart individual genetic effects, simultaneously estimating the phenotypic variation that is due to environment-persistent effects and due to genotype-environment interactions (GxE).

In genome-wide averages, we found that *cis* variation generally was more persistent to changes in environment, indicating that large parts of gene-environment interactions are are due to *trans* variation (*cis* GxE versus *trans* GxE) (see Figure 3a). Consequently, when grouping genes by annotated yeast pathways and ranking pathways by their average proportion of variance explained by GxE effects (the sum of *cis* GxE and *trans* GxE) (see Online Methods), we obtain processes that are enriched for *trans* variation (Figure 3b). These processes are plausibly implicated with the environmental stimulus (Supplementary Table 3). For example, Lysine biosynthesis (sce00300), consisting of 11 genes, has previously been found to be controlled by an interaction of genetic and environmental factors [31].

**Figure 3.**
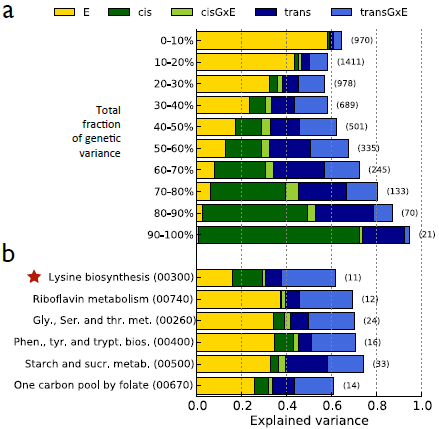
Variance component analysis of yeast gene expression levels in two environments [17]. For each gene, an LMM with variance components for local *cis* effects, genome-wide *trans* effects, environmental effects, and interaction effects between the environment and genetic variation (GxE) in *cis* as well as *trans* was fitted. **a**, Average variance component estimates for all genes grouped by the total proportion of variance explained by genetics (*cis* + *trans*). **b**, Average variance estimates for components of genes in the top six KEGG pathways with the largest average proportion of variance explained by GxE effects grouped by pathway membership; (*cis* GxE + *trans* GxE).

### Pathway-based genetic analysis across multiple environments

Next, to study how this variance components analysis complements multi-trait GWAS, we used multi-trait LMMs for association testing of the Lysine biosynthesis pathway, where we jointly considered all genes in the pathway (11) under both conditions (22 expression levels in total). Notably, this analysis shows how LIMIX can be used to integrate substantially more traits than previous applications of multi-trait LMMs [11, 12], and simultaneously spans both the axes of multiple genes in a pathway and multiple environments. First, we used LIMIX to construct a multi-trait single-locus model across all 22 traits. While other multi-trait methods, e.g. [12], can in principle be used to test for associations between single loci and this number of traits, we found LIMIX to greatly increase GWAS power (Optimally regularized model as in LIMIX versus an unregularized multi-trait LMM [12], Figure 4a, Supplementary Figure 6). This difference in power is because LIMIX allows for regularization of trait-trait covariances (Supplementary Figure 7, Online Methods), which we found to be critical for analyzing datasets with more than a handful of traits (Supplementary Figure 6; see also the Section below).

**Figure 4.**
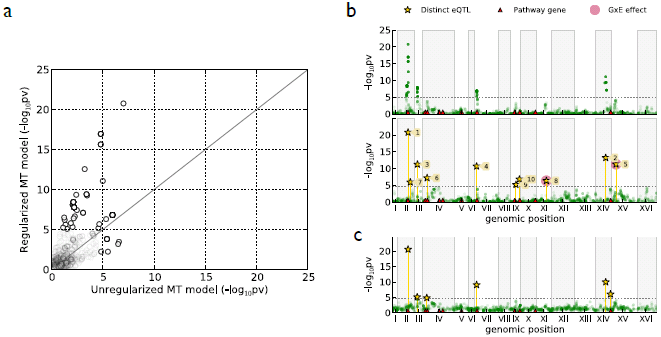
eQTL analysis of yeast gene expression in two environments [17], considering the Lysine biosynthesis pathway. **a**, Power comparison of alternative multi-trait LMMs on 11 genes in the pathway (22 measurements per sample). We considered an unregularized multi-trait LMM (*P*-values on the x-axes) versus the corresponding results obtained from an optimally regularized model (y-axes). Tests below the significance cutoff in both methods (Bonferroni adjusted *α* = 0.05) are shaded out. **b**, Joint multi-trait GWAS on the 11 genes in the pathway (22 measurements per sample). Upper panel: An any effect test finds 4 loci with significant associations (Bonferroni adjusted *α* = 0.05). Lower panel: A multi-locus multi-trait LMM with an any effect test identifies 10 SNPs with distinct genetic signals, each marked with a vertical bar. The *P*-value at iteration of insertion is marked by an asterisk. Two of the ten SNPs inserted also have a significant association in a joint test for gene-environment interaction effects (iterations 2 and 8 in the multi-locus model, see Supplementary Figure 8). **c**, A single-gene multi-locus LMMs, considering independent tests across genes but jointly across environments (pairwise multi-trait) identified 5 out of the 10 loci found using the multi-trait multi-locus model, and one additional association.

Next, turning our attention to the eQTL mapping results from this 22-degrees-of-freedom any effect test, the multi-trait LMM identified 4 significantly associated regions on chromosomes two, three, seven and fourteen (Bonferroni adjusted *α* = 0.05; see Figure 4b, upper panel). Analogous to the analysis on the NFBC data, we observed that multi-locus tests in combination with multi-trait modeling yielded an additional increase in detection power (10 distinct loci, 6 more than single-locus testing), suggesting complex genetic control of this pathway (Figure 4b, lower panel; Supplementary Table 4). Notably, we found that simpler pairwise multi-trait LMMs, where individual genes are tested independently but jointly across both environments (for example as in MTMM [11]), were underpowered (Figure 4c, Supplementary Table 4). The full multi-trait multi-locus model built in LIMIX uncovered five additional eQTLs that could not be detected by any other approach, 2 additional *cis* associations and 3 additional *trans* associations (Supplementary Table 4). This shows that multi-trait tests can add power not only for detecting *trans* effects, but also for detecting *cis* eQTLs, which are expected to target a single gene.

In addition to joint testing across all genes end environments, LIMIX allows to define application-specific tests to leverage the structure in the data, here phenotypic measurements that span both multiple genes in a pathway and two environmental conditions. To show this, we constructed a specific effect test for one of the two conditions (glucose), jointly testing for genotype-environment interactions across all genes (11 degrees-of-freedom test, Online Methods). In combination with the multi-locus model above, this test revealed GxE effects at two *trans* eQTLs, one on chromosome sixteen and one on chromosome fifteen (see Supplementary Figure 8c). The latter is located at *ATP19*, which encodes a subunit of a large enzyme complex (F1F0 ATP) required for ATP synthesis [32].

Finally, we repeated similar analyses on an even larger pathway (glycine, serine and threonine metabolism, sce00260, 24 genes), showing how LIMIX can be used for larger problems, and finding similar increases in mapping power (Supplementary Figure 9).

### Assessing and validating genetic models

The covariance models considered in these applications have been fit using out-of-sample prediction performance to adjust regularization and to guide model selection.

To validate this approach and to show that appropriately regularized models improve phenotype prediction on independent test data, we assessed the fitted models in a holdout experiment. First, we compared independent single-trait models (ST) to one joint multi-trait model (MT), re-estimating the parameters on each set of training data and computing best linear unbiased prediction (BLUP; see Online Methods) to predict the phenotypes of the test samples. On each dataset, we performed multiple repeats of cross validation, resulting in between 10 (human eQTL data) and 500 (NFBC) independent splits in training and test data (Online Methods). Without inclusion of fixed effects, the multi-trait model tended to show significant improvements in prediction over the single-trait model in all three application domains (Figure 5, Supplementary Figures 10-12). Next, we performed multi-locus modeling in each of the single-trait models and in the multi-trait model by running forward selection on the training data. Adding the loci detected in this way as fixed effects in the LMM consistently further improved the predictions over the corresponding model without fixed effects, both for independent BLUPs (ST multilocus versus ST) as well as for the joint multi-trait BLUP (MT multi-locus versus MT). The improved prediction performance suggests that the SNPs obtained from multi-locus modeling tag genuine genetic signal.

**Figure 5.**
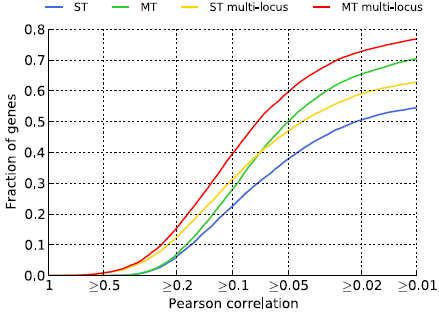
Prediction performance of alternative mixed models on the human eQTL dataset. Comparison of single-trait (ST) and multi-trait (MT) LMMs by out-of-sample prediction of isoform expression levels (all 9,246 genes, 10 fold cross validation). Multi-locus LMMs consistently achieved better prediction performance than the corresponding LMMs without fixed effects (ST multi-locus vs ST & MT multi-locus vs MT). On the whole, multi-trait LMMs predicted isoform levels more accurately than single-trait models. The combination of the multi-locus and multi-trait LMM achieved best prediction performance overall.

In the human eQTL study, we considered the largest number of independent prediction experiments, where for each of the 9,246 genes a separate experiment was carried out. When comparing predictions from a naive single-trait LMM (ST) to the multi-locus multi-trait LMM (MT multi-locus), the number of genes with correlation coefficient between predicted and true expression values of at least 0.5 increased from 1 to 73 and the number of genes with correlation coefficient of 0.1 or better went up from 2,370 to 3,438 (Figure 5). Notably, comparing individual contributions of multi-locus modeling and multi-trait modeling, we observed that for genes that could be accurately predicted (large *ρ*), multi-locus modeling was more important than multi-trait modeling, whereas genes that where hard to predict (low *ρ*) benefited mostly from multi-trait modeling (Figure 5). For a comparison of isoform predictions to predictions of gene-level expression estimates [33] and stratification by the number of isoforms or heritability, see Supplementary Figure 12).

## 3 Discussion

We have shown how a flexible mixed model framework allows to systematically extend LMMs to the genetic analysis of multiple related phenotypes. Whereas so far multi-trait analysis has been limited to GWAS on small groups of phenotypes [11, 12], due to computational complexity or overfitting, LIMIX can perform a much wider range of LMM applications on much larger numbers of phenotypes. We gave several example applications of LIMIX that address important questions about how genetics affect related phenotypes. These include association tests of different genetic designs, multi-locus analyses, variance components analysis, modeling of gene-environment interactions, and phenotype prediction.

In three different analyses we have demonstrated how multi-trait LMMs can be most effective in practice. Whereas multi-trait LMMs [12, 11] and multi-locus LMMs [22] have been proposed in isolation, we find that multi-trait LMMs are particularly powerful if combined with multilocus analyses. Owing to fast algorithms [23, 24, 12, 3, 4], it is now feasible to build complex models with a fairly large number of independent genetic effects that jointly affect a set of phenotypes in cohorts involving thousands of samples. In all cases these richer genetic models revealed extensive allelic heterogeneity [22].

Second, we have shown how multi-trait LMMs increase power and interpretation in genetic analyses of different sets of molecular phenotypes. As one example, we analyzed individual isoforms in human eQTL studies, where multi-trait analysis was used to dissect traditional gene-level genetic effects, revealing extensive isoform-specific genetic regulation. The use of LIMIX enabled to combine important concepts, such as accounting for expression heterogeneity [8, 9] with multi-trait modeling as well as tests involving multiple loci [22]. In a second example from yeast genetics, we have shown how multi-trait LMMs can be used to dissect genetic effects on genes in a pathway along two different axes, genes and environmental conditions. Our results show how genotype-environment interaction tests can be performed jointly for all genes in a pathway to increase power. Finally, we found that multi-trait multi-locus models improve genetic prediction of gene expression, a challenging task that has recently gained considerable attention [33].

LIMIX implements efficient algorithms that speed up multi-trait LMMs for analysis of larger numbers of traits [23, 24, 12]. Beyond the challenge of computational scalability, it is a well known problem in statistics that covariance estimation suffers from overfitting as the dimension of the trait-covariance increases [34], mainly because the parameters of these models grow quadratically whereas data only linearly.

Here, we address this problem by an automated model selection approach, which based on out-of-sample prediction tunes model complexity and regularization, thereby enabling the joint genetic analysis of even hundreds of traits (Online Methods). We have demonstrated the effectiveness of this approach in the yeast experiments, where we show that non-regularized models [12, 11] can lead to loss of power and poor prediction performance. The use of regularization to improve covariance estimation is well established in various different contexts [34, 23, 24], including its use for estimating genetic covariance matrices [35], however, to the best of our knowledge has not been used for multi-trait LMMs in GWAS. In our experiments we find that appropriate regularization of genetic models becomes increasingly important as the number of phenotype increases.

Finally, multi-trait LMMs should offer the greatest benefits if the phenotypes are correlated due to (partially) shared genetic causes or common environmental factors. Choosing the most effective analysis approach is non-trivial and as discussed in previous work [11, 12], an independent analysis of individual phenotypes can often be equally powered or even superior to complex modeling choices. While there is no silver bullet solution, LIMIX provides a wide range of models and approaches that can be objectively compared within the same framework. The in-depth applications of multi-trait modeling in the context of three application domains as presented here can serve as an example to guide future studies. Moreover, these applications show how the difficult task of selecting the most appropriate model can be addressed in a principled manner by out-of-sample prediction, a tool of tremendous value to determine model complexity and to validate complex genetic models.

## Online methods

### Multi-trait linear mixed models in LIMIX

Let *N* and *P* denote the number of samples and the number of traits in the genetic study. LIMIX models the *N* × *P* matrix of phenotype values as the sum of any number *J* of matrix-variate fixed effects ***F***_*j*_ and any number *I* of matrix-variate random effects ***U***_*i*_:

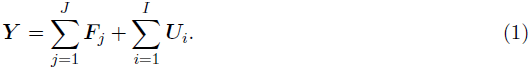

Here, fixed and random effects are denoted by ***F***_*j*_ and ***U***_*i*_ respectively. This formulation generalizes a wide range of previously proposed LMMs in genetics [1, 3, 12, 36, 37, 11, 22].

#### Fixed effects

Fixed effects represent known phenotypic covariates like gender, age, or ethnicity. In the case of a GWAS, additionally the SNP being tested is modeled as a fixed effect. For each sample *n* and trait *p* the *j*^th^ fixed effect *F*_*j*,*n*,*p*_ in (1) is given by the product of an effect size *β*_*j*_, a sample design *X*_*n*,*j*_ and a trait design *A*_*p*,*j*_:

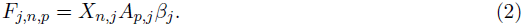

This fixed effect formulation generalizes the approach taken in single-trait LMMs, where fixed effects are represented as the product of a sample design matrix and the fitted effect size. The trait design matrix ***A*** is a design parameter that allows to specify alternative fixed effect designs in multiple trait LMMs (see multi-trait models below).

#### Random effects

Random effects are used to account for hidden influences such as a set of unknown causal variants, population structure or relatedness between samples. Even though these effects are unobserved, the distribution of the corresponding random effect can be described in terms of its covariance matrix. In LIMIX the covariance matrix of each random effect ***U***_*i*_ is assumed to factor into an *N*-by-*N* sample-to-sample covariance matrix ***R***_*i*_ and a *P*-by-*P* trait-to-trait covariance matrix ***C***_*i*_ as follows:

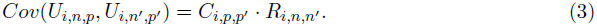

Again, this form generalizes random effects models in single-trait LMMs, where covariances are modeled between samples only.

LIMIX supports a flexible number of random effect terms in the model. For each random effect, both the trait and sample covariances can be chosen from a comprehensive class of covariance models, including models for population structure and relatedness [1, 3], models for hidden confounders inferred from high-dimensional phenotype data as in PANAMA [9], principal components models [38, 39], non-linear kernel functions [40], as well as combinations like sums and products of any of the aforementioned [41] (Supplementary Note 1). LIMIX performs efficient and robust parameter estimation and model checking for each of these covariance models.

#### Efficient estimation

In general, parameter estimation in multi-trait mixed models would be cubic in the number of values considered for ***Y***, yielding a runtime of *O*(*N*^3^*P*^3^). Due to the poor scalability this form of estimation is applicable only to small to medium-sized data sets. LIMIX utilizes specialized algorithms that, depending on the structure in the data, can speed up computation. For example in multi-trait analyses with up to two random effects the runtime of LIMIX is reduced to *O*(*N*^3^ + *P*^3^), whenever the matrix of phenotypes is fully observed [24, 12, 23, 42, 43] (Supplementary Note 1). For parameter inference, LIMIX uses a combination of gradient-based parameter inference (for random effects) and closed-form updates for fixed effects (Supplementary Note 1).

#### Statistical testing

By default LIMIX offers three different statistical tests for fixed effects using different trait design matrices, which are similar to those proposed in [11]. The *any effect test* is a *P* degrees-of-freedom test, where *P* is the number of traits, that tests whether there is at least one effect between the SNP and the phenotypes. The *common effect test* has 1 degree of freedom and tests if a SNP has the same effect size (and direction) across all the phenotypes. The *specific effect test* for trait *p* has 1 degree of freedom and tests if a SNP has a different effect on trait *p*.

Tests for designs different from the aforementioned ones can be performed by choosing appropriate trait design matrices both for the alternative and the null models.

#### Predictions

LIMIX implements best linear unbiased prediction (BLUP) [44, 45, 24] for phenotype prediction from the model in Equation (1), combining random effects and fitted fixed effects. Given the wealth of models made available in LIMIX, prediction in combination with cross-validation is indispensable for monitoring overfitting, model validation, and model selection (see also Cross-validation below). Moreover, the prediction tool from LIMIX can be used to impute missing phenotype values.

### Multi-trait models used in experiments

Here, we describe the different models built in LIMIX that have been used in this manuscript.

#### Variance decomposition

For the yeast eQTL analysis we jointly modeled the expression levels of each gene across *E* = 2 environments and used an LMM with an environment-specific mean modeled as fixed effect, two genetic random effect terms, accounting for contributions from proximal (*cis*) and distal (*trans*) genetic variation, and a non-genetic random effect term (noise)

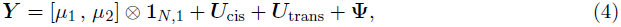

where *μ*_1_ and *μ*_2_ are environment-specific mean values. The variances of the random effects ***U***_cis_, ***U***_trans_, are each decomposed into an environment-persistent and environment specific variance component, whereas the noise *ψ* is fitted as free-form covariance

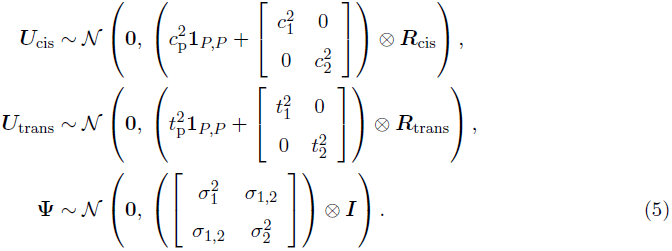

The samples-to-sample covariance matrices ***R***_cis_ and ***R***_trans_ are calculated as a genomic relatedness matrix estimated from the SNP data, where all SNPs within 100kb from the transcription start site where considered as *cis* and the remaining SNPs as *trans*. This model is similar to previous approaches that estimate persistent heritability values [46, 15] and builds on methods to estimate genetic variance components [5, 6]. We estimated the mean values *μ*_1_, and *μ*_2_, the environment persistent variances and covariances for *cis* variation 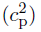, *trans* variation 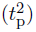, as well as the environment specific variances for all environments and the noise covariance parameters 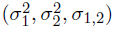, resulting in a total of 11 parameters, by maximum likelihood.

For each gene, the environment persistent variance component was reported as the variance of the block covariance (e.g. 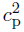 for *cis*) and environment specific variance components were reported as the average of the two corresponding environment-specific variances (e.g. 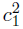 and 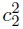 for *cis* GxE). The variance explained by direct environmental effects was estimated from the effect sizes of the fixed effects, *μ*_1_ and *μ*_2_ and the residual noise variance was estimated as the average of the noise variances (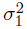 and 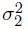). The overall variance components from both fixed and random effects were normalized to sum to 1 and subsequently averaged across the different sets of genes considered in Fig. 3.

For the pathway analysis, we considered only pathways for which at least 10 genes are contained in the Kegg pathway datbase [47], resulting in 77 pathways.

#### Genome-wide association studies

For multi-trait GWAS we used an LMM with trait-specific means modeled as fixed effects, a fixed effect containing the SNP being tested, a random effect accounting for genomic relatedness, and random noise:

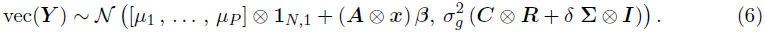

Here, vec(***Y***) denotes a stacking operation to creating an extended phenotpye vector from the phenotype matrix. This representation allows to express multivariate normal models as standard normal models with Kronecker-structured fixed effects and covariances; see Supplementary Note 1. The parameter *μ*_*p*_ denotes the mean for phenotype *p* ∈ (1,…, *P*) plus covariate effects, ***x*** denotes the SNP being tested, ***B*** is the effect size estimate for a model specified by the trait design matrix ***A***. The random effects are parameterized by the genomic relatedness matrix computed from genome-wide SNPs ***R***, the trait-to-trait genetic covariance matrix ***C***, the trait-to-trait noise covariance matrix **Σ**, and scaling factors 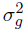 and *δ*. Statistical testing for models of form (6) is carried out by comparing the model likelihood under alternative designs ***A*** for alternative (foreground) model and null (background) model in a likelihood-ratio test. Testing a model without a SNP effect to a model with ***A*** = **1**_1,*P*_ in the alternative model results in a *common effect test*, i.e. a one degree of freedom test to detect associations with shared effect sizes and directions across phenotypes, comparing a null model with ***A*** = **1**_1,*P*_ to an alternative model with an additional degree of freedom results in a *specific effect test* to detect phenotype-specific regulation, and an *any effect test* with *P* degrees of freedom is achieved by comparing a null model without a SNP effect to an alternative model with ***A*** = ***I***_*P*_. Examples of more complex designs are considered for the pathway-based genetic analysis across environments. Here, a joint environment-specific effect test for *P* genes across two environments is achieved by comparing a model with ***A*** = ***I***_*P*_ ⊗ **1**_1,2_ to an alternative model with ***A*** = ***I***_*P*_ ⊗ ***I***_2_, resulting in a *P* degrees-of-freedom test.

For the transcript eQTL analysis in humans, the sample covariance matrix ***R*** was estimated estimated using the LIMIX-implementation of PANAMA [9], thereby accounting for hidden covariates and expression heterogeneity [29].

In all experiments, the trait-to-trait covariance matrices ***C*** and **Σ**, as well as the sample covariance matrix ***R***, have been estimated on the null model only. In genome-wide tests, the fitted matrices ***C*** and ***R*** were kept constant, however the scaling factors 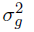 and *δ* were refit for each test, extending previous fast LMM algorithms [3, 4] to multi-trait analysis. More complex fits where all parameters are updated are possible as well [12], however at the price of additional computational cost.

#### Multi-locus models

We used LIMIX to implement **multi-trait multi-locus** LMMs, which combine concepts from the multi-locus LMM for single phenotypes [22] and single-locus LMMs for multiple traits [11]. Similar to [22], we employ forward selection however using a multi-trait LMM, iteratively including SNPs as fixed effect covariates into the model. In each iteration, the most significant SNP using the any effect test is included into the model as a covariate for the next iteration (we again consider the any effect design for each covariate). The **single-trait multi-locus** model considered in the analysis is the special case *P* = 1 of the general model. For experiments on the NFBC1966 data, we used the canonical GWAS significance threshold of *α* = 5 × 10^−8^, both for single-locus and multi-locus models. Similarly, for the genome-wide yeast eQTL mapping experiment, we employed a Bonferroni-adjusted significance level of *α* = 0.05. For the human eQTL study, association tests were carried out in a local *cis*-window around each gene (1mb up and downstream of genes) and significant eQTLs across genes were called at a nominal significance level of 1% FDR (estimated using Q-values [30]). For some analyses, we also considered a multi-trait multi-locus model using an alternative genetic design for forward selection. Similarly, in the yeast eQTL analysis we employed an interaction-test as forward-selection step (specific effect test for one of the two environmental conditions) to classify included SNPs as genotype-environment interactions (Figure 4b lower panel, Supplementary Figure 8).

#### Cross-validation

We validated the models used in each experiment by cross-validation, predicting phenotype values on holdout samples by BLUP. In prediction experiments, we considered a single-trait BLUP model (ST) and a multiple-trait BLUP (MT), performing joint predictions across phenotypes. Both models were extended by multi-locus approaches, where significant SNPs identified by the model were included as additional fixed effect terms (ST multi-locus & MT multi-locus). For the human eQTL analysis, prediction experiments were carried out using the genetic covariance, ignoring PANAMA [9] (see genome-wide association studies), as PANAMA random effects cannot be evaluated out-of-sample. For models that required tuning of regularization parameters we performed a separate round of cross-validation using the training data only to avoid in-sample estimation.

### Datasets and preprocessing

#### NFBC1966

We used results from the NFBC1966, consisting of phenotypic and genotypic data for 5,402 individuals [21]. We followed the normalization procedure described in [12], focusing on the four phenotypes CRP,LDL, HDL and TRIGL.

#### GEUVADIS eQTL dataset

We used the official release of genotype and raw RNA-Seq dataset from the GEUVAVIDS consortium [20]. First, raw RNA-Seq reads were mapped to the GRCh37.69 reference transcriptome using Bowtie [48], followed by isoform abundance estimation using BitSeq [27]. The MCMC samples of the BitSeq abundance estimates were processed in the natural log-scale, resulting in mean isoform abundance estimates for 39,211 genes. Considering only transcripts that were significantly expressed. We determined expression above background using a z-score cutoff, requiring that at least 5% of the samples were expressed 50% above the noise background (standard deviation). This filter led to the identification of 16,278 genes that express at least 1 transcript isoform. For the multi-trait analyses in Figure 2, only genes with 2-10 transcripts were used for analysis (9,246 genes). We ignored genes with more than 10 transcripts from the analysis (952 total), as the transcript reconstruction in BitSeq was found to be unstable for these genes [27]. For additional prediction experiments where gene-level abundance estimates were considered (Supplementary Figure 12), the full set of of 15,220 genes with at least one expressed transcript was used.

#### Yeast eQTL dataset

We used preprocessed gene expression and genotype data as provided by the authors [17]. The final dataset consisted of 109 individuals with 2,956 marker SNPs and 2 · 5,493 expression levels, profiled in glucose and ethanol growth media respectively. For the variance decomposition and multi-trait eQTL mapping, we considered Kegg pathways [47] with at least 10 annotated genes.

## Acknowledgements

The authors would like to thank Jennifer Listgarten, Leopold Parts, Andrew Brown and Amelie Baud for comments on the manuscript. The NFBC1966 Study is conducted and supported by the National Heart, Lung, and Blood Institute (NHLBI) in collaboration with the Broad Institute, UCLA, University of Oulu, and the National Institute for Health and Welfare in Finland. This manuscript was not prepared in collaboration with investigators of the NFBC1966 Study and does not necessarily reflect the opinions or views of the NFBC1966 Study Investigators, Broad Institute, UCLA, University of Oulu, National Institute for Health and Welfare in Finland and the NHLBI. O.S. was supported by a Marie Curie FP7 fellowship.

### Author contributions

C.L., F.P.C. & O.S. designed the study, developed methods, analysed data and wrote the paper. B. R. contributed to method development and wrote the paper.

### Competing financial interests

C. L. was employed at Microsoft while performing the research.

## Supplementary Information

### Supplementary Tables

1. **Supplementary Table 1** Tabular summary of significant associations in the NFBC datasets identified using alternative LMM methods.
2. **Supplementary Table 2** Tabular summary of *cis* associations identified in the Geuvadis dataset considering alternative multi-trait LMMs and tests.
3. **Supplementary table 3** Tabular summary of the average variance contribution in yeast pathways estimated using the variance decomposition model.
4. **Supplementary table 4** Tabular summary of significant associations on the *lysine bios*. yeast pathway.

### Supplementary Figures

1. **Supplementary Figure 1** *Q*–*Q*-plots for the multi-trait association tests on the NFBC dataset.
2. **Supplementary Figure 2** Sample covariance matrices used in mixed model analyses, either estimated from genetic kinship or using the PANAMA model in the human eQTL dataset.
3. **Supplementary Figure 3** Power comparison among different trait design tests on the human eQTL dataset.
4. **Supplementary Figure 4** Distribution of genomic control for the *cis* eQTL association analyses in the human eQTL dataset.
5. **Supplementary Figure 5** Distribution of the relative position of eQTLs identified using alternative multi-trait LMMs in the human eQTL dataset.
6. **Supplementary Figure 6** Power comparison of regularized and unregularized multi-trait LMMs for the analysis of the *lysine bios*. pathway in the yeast eQTL dataset.
7. **Supplementary Figure 7** Model selection approach to determine optimal regularization of the multi-trait LMM applied to the *lysine bios*. pathway in the yeast eQTL dataset.
8. **Supplementary Figure 8** Illustration of the step-wise inclusion of fixed effects used for multi-trait multi-locus LMMs applied to the *lysine bios*. pathway in the yeast eQTL dataset.
9. **Supplementary Figure 9** Multi-trait LMM analyses of 24 genes in the *glycine, s. and t. metabolism* pathway across two environments (48 traits) in the yeast eQTL dataset.
10. **Supplementary Figure 10** Assessment of alternative LMMs for out-of-sample prediction on the NFBC dataset.
11. **Supplementary Figure 11** Assessment of alternative LMMs for out-of-sample prediction of genes in the *lysine bios*. pathway in the yeast eQTL dataset.
12. **Supplementary Figure 12** Assessment of alternative LMMs for out-of-sample prediction of isoform and gene expression levels in the human eQTL dataset.

### Supplementary Notes

1. **Supplementary Note 1:** Supplementary Methods and additional theory.

