## Supplementary material for "LIMIX: genetic analysis of multiple traits": Data Supplement 1 (from another version of this preprint: version 1, 2014-05-20)

### Supplemental Information
### Multivariate analysis of heritable traits

authors

February 28, 2014

### 1 Limix methods and implementation

Limix is a flexible mixed model framework, allowing to efficiently formulate a variety of different mixed model analyses. Complex covariance functions that describe random effects can be combined from building blocks, using addition and multiplication as basic operations (Section 1.1). Inference in these models can be performed by gradient-based optimisation of the marginal likelihood or a regularized version thereof (Section 1.2). Section 1.4 describes how the same framework can be extended to perform efficient inference in the special case of matrix variate data and Section 1.5 outlines the software implementation and exemplifies the basic usage.

In the most general formulation, we assume that the data $\boldsymbol{y}$ is multivariate normal distribution with mean $\boldsymbol{\mu}$ and covariance matrix $\boldsymbol{\Sigma}$

$$\boldsymbol{y} \sim \mathcal{N}\left(\boldsymbol{\mu}_{\boldsymbol{\theta}}\,;\,\boldsymbol{\Sigma}_{\boldsymbol{\theta}}\right), \tag{1.1}$$

where $\boldsymbol{\theta}$ is the set of model parameters. In linear mixed models $\boldsymbol{\mu}_{\theta}$ typically is a linear function $\boldsymbol{\mu}_{\theta} = \boldsymbol{X}\boldsymbol{\beta}$, parametrized by a fixed design matrix $\boldsymbol{X}$ and fixed-effect parameters $\boldsymbol{\beta}$. The covariance matrix $\boldsymbol{\Sigma}_{\theta}$ is parametrized by individual variance components or covariance functions, corresponding to random effects in the model. In this instance, the set of model parameters $\boldsymbol{\theta}$ is given by the union of the fixed effects $\boldsymbol{\beta}$ and the variance parameters.

#### 1.1 Kernel methods for defining composite covariance functions

Defining property of a valid covariance matrix is a positive semi-definite symmetric matrix. In Limix we utilize a key insight from the kernel machines literature that allows construction of covariance matrices by combing a set of covariance matrices. First, we define a set of base covariance functions used in Limix (Section 1.1.1). In Section 1.1.2, we then define basic operations to combine two base covariance matrices $\boldsymbol{A}$ and $\boldsymbol{B}$ using sum and multiplication operators, which obey positive semi-definiteness of the resulting matrix $\boldsymbol{C}$. By iteratively applying these operations we can obtain complex covariance matrices from simpler base covariance models.

##### 1.1.1 Base covariance models

Limix provides implementations for a range of covariance matrices, where the most important examples are defined here. A more rigorous treatment of covariance functions including the definition of more extensive examples can be found in Rasmussen and Williams [2005]. Each covariance is parameterised by a set of parameters $\boldsymbol{\theta}$, giving rise to a semi-positive definite covariance matrix $\boldsymbol{C}(\boldsymbol{\theta})$. For inference, the matrix derivatives of $\boldsymbol{C}$ w.r.t $\boldsymbol{\theta}_i$ are needed, see also Section 1.2.

###### Fixed covariance matrix

If the covariance $\boldsymbol{A}$ between the samples is observed, we have to learn the amplitude $\sigma_g^2 > 0$:

$$\boldsymbol{C}(\boldsymbol{\theta}) = \sigma_g^2 \boldsymbol{A}, \quad \frac{\partial \boldsymbol{C}}{\partial \sigma_g^2} = \boldsymbol{A} \tag{1.2}$$

Here, $\boldsymbol{\theta} = [\sigma_g^2]$.

###### Linear covariance function

The linear covariance function can be derived by marginalizing out the weights in a random effect model $\boldsymbol{X}\boldsymbol{\beta}$, assuming $\boldsymbol{\beta} \sim \mathcal{N}(0, \sigma_g^2)$. This results in a covariance of the form $\boldsymbol{C}(\boldsymbol{\theta}) = \sigma_g^2 \boldsymbol{X}\boldsymbol{X}^\top$, where again $\boldsymbol{\theta} = [\sigma_g^2]$. This covariance can be used to model polygenic effects if $\boldsymbol{X}$ corresponds to genome-wide SNP data.

###### Free form covariance model

In practice, the covariance between phenotypes is not observed and hence it's parameters need to be inferred from data. The most general form is the free form covariance, which is a general $N \times N$ matrix that obeys the semi positive-definiteness constraints. To this end, we parametrize the covariance using cholesky factors, as any semi-positive definite matrix has a valid cholesky decomposition.

$$\boldsymbol{C}(\boldsymbol{\theta}) = \boldsymbol{L}\boldsymbol{L}^\top, \quad \frac{\partial \boldsymbol{C}}{\partial \boldsymbol{L}} = \boldsymbol{L}\partial(\boldsymbol{L})^\top + \partial(\boldsymbol{L})\boldsymbol{L}^\top \tag{1.3}$$

where $\boldsymbol{L}$ is a lower triangular matrix of covariance parameters, i.e. $\boldsymbol{\theta} = [\boldsymbol{L}_{0,0}, \boldsymbol{L}_{1,0}, \boldsymbol{L}_{1,1}, \ldots, \boldsymbol{L}_{0,N}, \boldsymbol{L}_{N,N}]$. The gradient with respect to a matrix field $\boldsymbol{L_{ij}}$ can be obtained by applying the chain rule $\frac{\partial \boldsymbol{C}}{\partial \boldsymbol{L_{ij}}} = \frac{\partial \boldsymbol{C}}{\partial \boldsymbol{L}} \frac{\partial \boldsymbol{L}}{\partial \boldsymbol{L_{ij}}}$.

###### Low-rank covariance model

We can restrict the phenotypes to lie in a lower-dimensional linear subspace, by using a low-rank parameterization of the covariance matrix

$$\boldsymbol{C}(\boldsymbol{\theta}) = \boldsymbol{Z}\boldsymbol{Z}^\top, \quad \frac{\partial \boldsymbol{C}}{\partial \boldsymbol{Z}} = 2\boldsymbol{Z} \tag{1.4}$$

where $\boldsymbol{Z}$ is a $N \times K$ matrix, with $K < N$, and the parameters are its entries, i.e. $\boldsymbol{\theta} = [\boldsymbol{Z}_{0,0}, \boldsymbol{Z}_{0,1}, \ldots, \boldsymbol{Z}_{0,K}, \ldots, \boldsymbol{Z}_{n,0}, \ldots, \boldsymbol{Z}_{N,K}]$. The gradient with respect to an entry $\boldsymbol{Z}_{ij}$ can then again be obtained by using the chain rule $\frac{\partial \boldsymbol{C}}{\partial \boldsymbol{Z}_{ij}} = \frac{\partial \boldsymbol{C}}{\partial \boldsymbol{Z}} \frac{\partial \boldsymbol{Z}}{\partial \boldsymbol{Z}_{ij}}$.

###### Diagonal covariance matrix

The diagonal covariance matrix is often used to model the noise between the phenotypes. By restricting the matrix to be diagonal, we implicitly assume that the noise is independently, but not identically distributed

$$\boldsymbol{C}(\boldsymbol{d}) = \operatorname{diag}(\boldsymbol{d}), \quad \frac{\partial \boldsymbol{C}}{\partial \boldsymbol{d}} = \boldsymbol{I}, \tag{1.5}$$

where $\boldsymbol{\theta} = [\boldsymbol{d}_1, \ldots, \boldsymbol{d}_N]$.

##### 1.1.2 Composite covariance functions

First, the sum of two covariance matrices is a covariance matrix.

$$\boldsymbol{A} + \boldsymbol{B} = \boldsymbol{C}, \tag{1.6}$$

Additive combinations of covariance matrices allow for variance decomposition and can be interpreted as multiple independent random effects.Second, the point-wise product of two covariance matrices is a valid covariance matrix,

$$\boldsymbol{A} \odot \boldsymbol{B} = \boldsymbol{C} \tag{1.7}$$

and the Kronecker product of two covariances matrices is as well (see also Section 1.4)

$$\boldsymbol{A} \otimes \boldsymbol{B} = \boldsymbol{C}. \tag{1.8}$$

The gradient of the composite covariance functions can be obtained by applying iteratively the sum and product rule TODO $\partial$ is now a gradient, not derivative.

$$\frac{\partial}{\partial \boldsymbol{\theta}}(\boldsymbol{A}+\boldsymbol{B}) = \frac{\partial}{\partial \boldsymbol{\theta}}\boldsymbol{A} + \frac{\partial}{\partial \boldsymbol{\theta}}\boldsymbol{B} \tag{1.9}$$

$$\frac{\partial}{\partial \boldsymbol{\theta}}(\boldsymbol{A}\odot\boldsymbol{B}) = \frac{\partial}{\partial \boldsymbol{\theta}}(\boldsymbol{A})\odot\boldsymbol{B} + \boldsymbol{A}\odot\frac{\partial}{\partial \boldsymbol{\theta}}(\boldsymbol{B}) \tag{1.10}$$

$$\frac{\partial}{\partial \boldsymbol{\theta}}(\boldsymbol{A}\otimes\boldsymbol{B}) = \frac{\partial}{\partial \boldsymbol{\theta}}(\boldsymbol{A})\otimes\boldsymbol{B} + \boldsymbol{A}\otimes\frac{\partial}{\partial \boldsymbol{\theta}}(\boldsymbol{B}). \tag{1.11}$$

#### 1.2 Inference and parameter estimation

##### 1.2.1 Maximizing the model likelihood

We perform parameter estimation by maximizing the log likelihood. The gradient of the log likelihood is optimized using LBFGS [Liu and Nocedal, 1989].

$$\begin{aligned}\frac{\partial}{\partial \boldsymbol{\theta}_i}\log\mathcal{N}\left(\boldsymbol{y}\,\middle|\,\boldsymbol{\mu_\theta}\,;\,\boldsymbol{\Sigma_\theta}\right) = &-\frac{1}{2}\mathrm{tr}\left(\boldsymbol{\Sigma}_{\boldsymbol{\theta}}^{-1}\frac{\partial\boldsymbol{\Sigma_\theta}}{\partial\boldsymbol{\theta}_i}\right) + \frac{1}{2}(\boldsymbol{y}-\boldsymbol{\mu_\theta})^\top\boldsymbol{\Sigma}_{\boldsymbol{\theta}}^{-1}\frac{\partial\boldsymbol{\Sigma_\theta}}{\partial\boldsymbol{\theta}_i}\boldsymbol{\Sigma}_{\boldsymbol{\theta}}^{-1}(\boldsymbol{y}-\boldsymbol{\mu_\theta}) \\ &-(\boldsymbol{y}-\boldsymbol{\mu_\theta})^\top\boldsymbol{\Sigma}_{\boldsymbol{\theta}}^{-1}\frac{\partial(\boldsymbol{y}-\boldsymbol{\mu_\theta})}{\partial\boldsymbol{\theta}_i}\end{aligned} \tag{1.12}$$

##### 1.2.2 Regularization

Estimating the covariance matrix between phenotypes is hindered by the large parameter space: for free-form covariance matrices, the number of parameters is growing quadratically with the number of phenotypes, while the number of new data points is only growing linearly. If not accounted for, this will lead to overfitting, i.e. the model does not only explain the signal of the data but also the noise, which in turn leads to a bad generalization behavior and a loss of interpretability. A natural way to prevent this is to add a regularization term over the covariance matrix

$$\min_{\boldsymbol{\theta}} \underbrace{-\log\mathcal{N}\left(\boldsymbol{y}\,\middle|\,\boldsymbol{\mu_\theta}\,;\,\boldsymbol{\Sigma_\theta}\right)}_{\text{data fit}} + \underbrace{\mathrm{Reg}(\boldsymbol{\Sigma_\theta})}_{\text{regularizer}}. \tag{1.13}$$

While the first term maximizes the fit between the data and the model, the second term acts as a penalizer that prevents the model from becoming too complex. In practice, we used the sum-of-squares penalty on the off-diagonal entries as regularization function $\mathrm{Reg}(\boldsymbol{\Sigma_\theta}) = \lambda\sum_{i\neq j}{(\boldsymbol{\Sigma_\theta})_{ij}}^2$, where $\lambda$ is the trade-off parameter between fitting the data and regularizing. We do not penalize the diagonal elements, to retain an unbiased estimate of the heritabilities. From the Bayesian perspective, the regularization term is equivalent to an isotropic Gaussian prior, with zero mean.

**Out of sample prediction** In practice, it is often hard to find the right trade-off between model fit and regularization. We used out of sample prediction accuracy to find the appropriate $\lambda$. Predictions for new data points can be carried out by conditioning on the observed data

$$\begin{aligned}p(\boldsymbol{y}^*|\boldsymbol{y},\boldsymbol{\theta}) = &\mathcal{N}\left(\boldsymbol{y}^*|\boldsymbol{\mu_\theta}(\boldsymbol{X}^*) + \boldsymbol{\Sigma_\theta}(\boldsymbol{X}^*,\boldsymbol{X})\boldsymbol{\Sigma}_{\boldsymbol{\theta}}^{-1}(\boldsymbol{y}-\boldsymbol{\mu_\theta});\right. \\ &\left.\boldsymbol{\Sigma_\theta}(\boldsymbol{X}^*,\boldsymbol{X}^*) - \boldsymbol{\Sigma_\theta}(\boldsymbol{X}^*,\boldsymbol{X})\boldsymbol{\Sigma}_{\boldsymbol{\theta}}^{-1}\boldsymbol{\Sigma_\theta}(\boldsymbol{X},\boldsymbol{X})^*\right),\end{aligned} \tag{1.14}$$

where $\boldsymbol{\mu_\theta}(\boldsymbol{X}^*)$ is the mean function on the test inputs, $\boldsymbol{\Sigma_\theta}(\boldsymbol{X},\boldsymbol{X})^*$ depicts the covariance between the training and the test samples, and $\boldsymbol{\Sigma_\theta}(\boldsymbol{X}^*,\boldsymbol{X}^*)$ between the test samples.

#### 1.3 Kronecker product identities

Let $\boldsymbol{A}$ be a $N \times M$ matrix, and $\boldsymbol{B}$ be a $P \times Q$ matrix. The Kronecker product $\boldsymbol{A} \otimes \boldsymbol{B}$ is a $NP \times MQ$ matrix and defined as follows

$$\boldsymbol{A} \otimes \boldsymbol{B} = \begin{pmatrix} A_{11}\boldsymbol{B} & \dots & A_{1n}\boldsymbol{B} \\ \vdots & \ddots & \vdots \\ A_{m1}\boldsymbol{B} & \dots & A_{mn}\boldsymbol{B} \end{pmatrix} \tag{1.15}$$

Due to its block structure, the Kronecker product has a number of nice properties that speed up inference[Bernstein, 2009, Petersen and Pedersen, 2006]. In particular, the dot product between two Kronecker products is a Kronecker product again

$$(\boldsymbol{A} \otimes \boldsymbol{B})(\boldsymbol{C} \otimes \boldsymbol{D}) = \boldsymbol{AC} \otimes \boldsymbol{BD}, \tag{1.16}$$

where $\boldsymbol{C} \in \mathbb{R}^{M \times R}$ and $\boldsymbol{D} \in \mathbb{R}^{Q \times S}$. It can be evaluated in $O(NM^2R + PQ^2S)$ time, while the naive runtime is $O(NPM^2Q^2RS)$.

Let vec be an operation that concatenates the columns of a $N \times M$ matrix into a vector of length $N \cdot M$. The product of Kronecker product and a vectorized matrix can be computed efficiently

$$(\boldsymbol{A} \otimes \boldsymbol{B}) \operatorname{vec}(\boldsymbol{Y}) = \operatorname{vec}\left(\boldsymbol{BYA}^\top\right) \tag{1.17}$$

bringing the runtime down to $O(PQ^2N + QN^2M)$ from $O(NPMQ)$.

Let $\boldsymbol{U_A S_A U_A^\top}$ be the eigenvalue decomposition of $\boldsymbol{A} \in \mathbb{R}^{N \times N}$ and $\boldsymbol{U_B S_B U_B^\top}$ the eigenvalue decomposition of $\boldsymbol{B} \in \mathbb{R}^{P \times P}$. The eigenvalue decomposition of the Kronecker product $\boldsymbol{A} \otimes \boldsymbol{B}$ can be synthesized by the eigenvalue decompositions of its components

$$\boldsymbol{A} \otimes \boldsymbol{B} = (\boldsymbol{U_A} \otimes \boldsymbol{U_B})(\boldsymbol{S_A} \otimes \boldsymbol{S_B})\left(\boldsymbol{U_A}^\top \otimes \boldsymbol{U_B}^\top\right) \tag{1.18}$$

leading to a runtime reduction from $O(N^3P^3)$ to $O(N^3 + P^3)$.

#### 1.4 Efficient inference for matrix variate mixed models

In Section 1.1, we gave a general overview over existing covariance functions. We now consider the special case in which the covariance matrix can be written as a sum of two Kronecker products and the fixed design matrix also exhibits Kronecker structure

$$\mathcal{N}\left(\operatorname{vec}(\boldsymbol{Y}) \,\middle|\, \sum_{j=1}^{J} (\boldsymbol{A}_j \otimes \boldsymbol{X}_j) \operatorname{vec}(\boldsymbol{B}_j)\,;\, \underbrace{\boldsymbol{C}(\boldsymbol{\theta_C}) \otimes \boldsymbol{R}(\boldsymbol{\theta_R})}_{\text{signal}} + \underbrace{\boldsymbol{\Sigma}(\boldsymbol{\theta_\Sigma}) \otimes \boldsymbol{I}}_{\text{noise}}\right), \tag{1.19}$$

Here, $\boldsymbol{Y}$ denotes the $N \times P$ data matrix with $N$ samples and $P$ phenotypes. For the fixed effect $j$, the matrix $\boldsymbol{B}_j \in \mathbb{R}^{M_j \times D_j}$ denotes the fixed-effects, $\boldsymbol{A}_j \in \mathbb{R}^{P \times M_j}$ is the design matrix of the column effects and $\boldsymbol{X}_j \in \mathbb{R}^{N \times D_j}$ of the row effects. We define $\boldsymbol{R}(\boldsymbol{\theta_R})$ as the signal row covariance of the data matrix and $\boldsymbol{C}(\boldsymbol{\theta_C})$ as the signal column covariance matrix. Similarly, the noise column covariance matrix is given by $\boldsymbol{\Sigma}(\boldsymbol{\theta_\Sigma})$. For the sake of clarity, we further assume that the noise row covariance matrix is the identity $\boldsymbol{I}$, the extension to an arbitrary noise row covariance matrix is straightforward. For notational convenience, we will also drop the dependence of the covariance matrices on additional hyperparameters from now on.

The merits of this class of models lie in its tractability: as we show in the following, efficient inference can be performed in $O(N^3 + P^3)$ time and in $O(N^2 + P^2)$ space, whereas an arbitrary covariance matrix of size $NP$ has runtime $O(N^3P^3)$ and a memory requirement of $O(N^2P^2)$.

##### 1.4.1 Log likelihood evaluation

The log likelihood of the data is given by:

$$\log \mathcal{L} = -\frac{NP}{2}\log(2\pi) - \frac{1}{2}\log|\boldsymbol{C}\otimes\boldsymbol{R}+\boldsymbol{\Sigma}\otimes\boldsymbol{I}| - \frac{1}{2}\text{vec}(\boldsymbol{Y}_r)^\top(\boldsymbol{C}\otimes\boldsymbol{R}+\boldsymbol{\Sigma}\otimes\boldsymbol{I})^{-1}\text{vec}(\boldsymbol{Y}_r), \tag{1.20}$$

where $\boldsymbol{Y}_r$ is the residual phenotype, after the fixed effects have been subtracted from the data:

$$\begin{aligned}\text{vec}(\boldsymbol{Y}_r) &= \text{vec}(\boldsymbol{Y}) - \sum_{j=1}^{J}(\boldsymbol{A}_j\otimes\boldsymbol{X}_j)\,\text{vec}(\boldsymbol{B}_j)\\ &= \text{vec}\left(\boldsymbol{Y} - \sum_{j=1}^{J}\boldsymbol{X}_j\boldsymbol{B}_j\boldsymbol{A}_j^\top\right)\end{aligned} \tag{1.21}$$

Let $\boldsymbol{U_\Sigma S_\Sigma U_\Sigma}^\top$ be the eigenvalue decomposition of $\boldsymbol{\Sigma}$. As shown in [Stegle et al., 2011a, Rakitsch et al., 2013], we can whiten the covariance matrix in a two-step procedure. First, we whiten the noise by factoring it out. Second, we exploit the fact that the eigenvalue decomposition of a Kronecker product is compatible with a constant diagonal matrix to whiten the remaining covariance matrix:

$$\begin{aligned}\boldsymbol{K} &= \boldsymbol{C}\otimes\boldsymbol{R}+\boldsymbol{\Sigma}\otimes\boldsymbol{I}\\
&\overset{(1.18)}{=} \boldsymbol{C}\otimes\boldsymbol{R}+\boldsymbol{U_\Sigma S_\Sigma U_\Sigma^\top}\otimes\boldsymbol{I}\\
&\overset{(1.16)}{=} \left(\boldsymbol{U_\Sigma S_\Sigma}^{\frac{1}{2}}\otimes\boldsymbol{I}\right)\left(\underbrace{\boldsymbol{S_\Sigma}^{-\frac{1}{2}}\boldsymbol{U_\Sigma}^\top\boldsymbol{C}\boldsymbol{U_\Sigma}\boldsymbol{S_\Sigma}^{-\frac{1}{2}}}_{\tilde{\boldsymbol{C}}}\otimes\boldsymbol{R}+\boldsymbol{I}\otimes\boldsymbol{I}\right)\left(\boldsymbol{S_\Sigma}^{\frac{1}{2}}\boldsymbol{U_\Sigma}^\top\otimes\boldsymbol{I}\right)\\
&\overset{(1.18)}{=} \left(\boldsymbol{U_\Sigma S_\Sigma}^{\frac{1}{2}}\otimes\boldsymbol{I}\right)\left(\boldsymbol{U_{\tilde{C}}S_{\tilde{C}}U_{\tilde{C}}^\top}\otimes\boldsymbol{U_R S_R U_R^\top}+\boldsymbol{I}\otimes\boldsymbol{I}\right)\left(\boldsymbol{S_\Sigma}^{\frac{1}{2}}\boldsymbol{U_\Sigma}^\top\otimes\boldsymbol{I}\right)\\
&\overset{(1.16)}{=} \left(\boldsymbol{U_\Sigma S_\Sigma}^{\frac{1}{2}}\boldsymbol{U_{\tilde{C}}}\otimes\boldsymbol{U_R}\right)\left(\boldsymbol{S_{\tilde{C}}}\otimes\boldsymbol{S_R}+\boldsymbol{I}\otimes\boldsymbol{I}\right)\left(\boldsymbol{U_{\tilde{C}}^\top}\boldsymbol{S_\Sigma}^{\frac{1}{2}}\boldsymbol{U_\Sigma}^\top\otimes\boldsymbol{U_R^\top}\right),\end{aligned} \tag{1.22}$$

where $\boldsymbol{U_{\tilde{C}}S_{\tilde{C}}U_{\tilde{C}}^\top}$ is the eigenvalue decomposition of $\tilde{\boldsymbol{C}}$. The log determinant can then be written as

$$\log|\boldsymbol{C}\otimes\boldsymbol{R}+\boldsymbol{\Sigma}\otimes\boldsymbol{I}| \overset{|AB|=|A|\cdot|B|}{=} \log|\boldsymbol{S_\Sigma}\otimes\boldsymbol{I}| + \log|\boldsymbol{S_{\tilde{C}}}\otimes\boldsymbol{S_R}+\boldsymbol{I}\otimes\boldsymbol{I}| \tag{1.23}$$

$$\overset{|A\otimes B|=|A|^P\cdot|B|^N}{=} N\sum_{p=1}^{P}\log\boldsymbol{S_\Sigma}[p,p] + \sum_{p=1}^{P}\sum_{n=1}^{N}\log\left(\boldsymbol{S_{\tilde{C}}}[p,p]\boldsymbol{S_R}[n,n]+1\right) \tag{1.24}$$

We can evaluate the squared form efficiently as follows

$$\begin{aligned}&\text{vec}(\boldsymbol{Y}_r)^\top(\boldsymbol{C}\otimes\boldsymbol{R}+\boldsymbol{\Sigma}\otimes\boldsymbol{I})^{-1}\text{vec}(\boldsymbol{Y}_r)\\
\overset{(1.22)}{=}\;&\text{vec}(\boldsymbol{Y}_r)^\top\left[\left(\boldsymbol{U_\Sigma S_\Sigma}^{\frac{1}{2}}\boldsymbol{U_{\tilde{C}}}\otimes\boldsymbol{U_R}\right)\left(\boldsymbol{S_{\tilde{C}}}\otimes\boldsymbol{S_R}+\boldsymbol{I}\otimes\boldsymbol{I}\right)\left(\boldsymbol{U_{\tilde{C}}^\top}\boldsymbol{S_\Sigma}^{\frac{1}{2}}\boldsymbol{U_\Sigma}^\top\otimes\boldsymbol{U_R^\top}\right)\right]^{-1}\text{vec}(\boldsymbol{Y}_r)\\
=\;&\left[\left(\boldsymbol{U_\Sigma S_\Sigma}^{\frac{1}{2}}\boldsymbol{U_{\tilde{C}}}\otimes\boldsymbol{U_R}\right)^{-1}\text{vec}(\boldsymbol{Y}_r)\right]^\top\left(\boldsymbol{S_{\tilde{C}}}\otimes\boldsymbol{S_R}+\boldsymbol{I}\otimes\boldsymbol{I}\right)^{-1}\left[\left(\boldsymbol{U_\Sigma S_\Sigma}^{\frac{1}{2}}\boldsymbol{U_{\tilde{C}}}\otimes\boldsymbol{U_R}\right)^{-1}\text{vec}(\boldsymbol{Y}_r)\right]\\
=\;&\left[\left(\boldsymbol{U_{\tilde{C}}^\top}\boldsymbol{S_\Sigma}^{-\frac{1}{2}}\boldsymbol{U_\Sigma^\top}\otimes\boldsymbol{U_R^\top}\right)\text{vec}(\boldsymbol{Y}_r)\right]^\top\left(\boldsymbol{S_{\tilde{C}}}\otimes\boldsymbol{S_R}+\boldsymbol{I}\otimes\boldsymbol{I}\right)^{-1}\left[\left(\boldsymbol{U_{\tilde{C}}^\top}\boldsymbol{S_\Sigma}^{-\frac{1}{2}}\boldsymbol{U_\Sigma^\top}\otimes\boldsymbol{U_R^\top}\right)\text{vec}(\boldsymbol{Y}_r)\right]\\
\overset{(1.17)}{=}\;&\text{vec}\left[\boldsymbol{U_R^\top}\boldsymbol{Y}_r\boldsymbol{U_\Sigma}\boldsymbol{S_\Sigma}^{-\frac{1}{2}}\boldsymbol{U_{\tilde{C}}}\right]^\top\left(\boldsymbol{S_{\tilde{C}}}\otimes\boldsymbol{S_R}+\boldsymbol{I}\otimes\boldsymbol{I}\right)^{-1}\text{vec}\left[\boldsymbol{U_R^\top}\boldsymbol{Y}_r\boldsymbol{U_\Sigma}\boldsymbol{S_\Sigma}^{-\frac{1}{2}}\boldsymbol{U_{\tilde{C}}}\right]\\
=\;&\text{vec}\left[\boldsymbol{U_R^\top}\boldsymbol{Y}_r\boldsymbol{U_\Sigma}\boldsymbol{S_\Sigma}^{-\frac{1}{2}}\boldsymbol{U_{\tilde{C}}}\right]^\top\text{vec}\left[\boldsymbol{D}\odot\boldsymbol{U_R^\top}\boldsymbol{Y}_r\boldsymbol{U_\Sigma}\boldsymbol{S_\Sigma}^{-\frac{1}{2}}\boldsymbol{U_{\tilde{C}}}\right],\end{aligned} \tag{1.25}$$

where $\boldsymbol{D}$ is a $N\times P$ matrix defined by the entries

$$\boldsymbol{D}[n,p] = \frac{1}{\boldsymbol{S_{\tilde{C}}}[p,p]\otimes\boldsymbol{S_R}[n,n]+1}. \tag{1.26}$$

The inverse of the covariance term times a vector is therewith:

$$(\boldsymbol{C} \otimes \boldsymbol{R} + \boldsymbol{\Sigma} \otimes \boldsymbol{I})^{-1} \operatorname{vec}(\boldsymbol{Y}_r) = \left(\boldsymbol{U_\Sigma} \boldsymbol{S_\Sigma}^{-\frac{1}{2}} \boldsymbol{U_{\tilde{C}}} \otimes \boldsymbol{U_R}\right) \operatorname{vec}\left[\underbrace{\boldsymbol{D} \odot \boldsymbol{U_R}^\top \boldsymbol{Y}_r \boldsymbol{U_\Sigma} \boldsymbol{S_\Sigma}^{-\frac{1}{2}} \boldsymbol{U_{\tilde{C}}}}_{\tilde{\boldsymbol{Y}}_r}\right] \tag{1.27}$$

#### 1.4.2 Estimation of the covariance parameters

We want to evaluate the gradient with respect to a particular covariance parameter $\theta \in \{\boldsymbol{\theta_C}, \boldsymbol{\theta_R}, \boldsymbol{\theta_\Sigma}\}$. The derivative consists of the derivative of the determinant term and the derivative of the squared form. Each of these is given separately for the row covariance parameter $\theta_r \in \boldsymbol{\theta_R}$. The gradients for the column covariance parameters can be derived analogously and are omitted for brevity.

We start with the gradient of the log determinant:

$$\begin{aligned}
& \frac{\partial}{\partial \theta_r} \log |\boldsymbol{C} \otimes \boldsymbol{R} + \boldsymbol{\Sigma} \otimes \boldsymbol{I}| \\
= \quad & \operatorname{tr}\left((\boldsymbol{C} \otimes \boldsymbol{R} + \boldsymbol{\Sigma} \otimes \boldsymbol{I})^{-1} \left(\boldsymbol{C} \otimes \frac{\partial}{\partial \theta_r} \boldsymbol{R}\right)\right) \\
\overset{(1.22)}{=} \quad & \operatorname{tr}\left(\left[\left(\boldsymbol{U_\Sigma} \boldsymbol{S_\Sigma}^{\frac{1}{2}} \boldsymbol{U_{\tilde{C}}} \otimes \boldsymbol{U_R}\right) (\boldsymbol{S_{\tilde{C}}} \otimes \boldsymbol{S_R} + \boldsymbol{I} \otimes \boldsymbol{I}) \left(\boldsymbol{U_{\tilde{C}}}^\top \boldsymbol{S_\Sigma}^{\frac{1}{2}} \boldsymbol{U_\Sigma}^\top \otimes \boldsymbol{U_R}^\top\right)\right]^{-1} \left(\boldsymbol{C} \otimes \frac{\partial}{\partial \theta_r} \boldsymbol{R}\right)\right) \\
\overset{(AB)^{-1}=B^{-1}A^{-1}}{=} \quad & \operatorname{tr}\left(\left(\boldsymbol{U_{\tilde{C}}}^\top \boldsymbol{S_\Sigma}^{\frac{1}{2}} \boldsymbol{U_\Sigma}^\top \otimes \boldsymbol{U_R}^\top\right)^{-1} (\boldsymbol{S_{\tilde{C}}} \otimes \boldsymbol{S_R} + \boldsymbol{I} \otimes \boldsymbol{I})^{-1} \left(\boldsymbol{U_\Sigma} \boldsymbol{S_\Sigma}^{\frac{1}{2}} \boldsymbol{U_{\tilde{C}}} \otimes \boldsymbol{U_R}\right)^{-1} \left(\boldsymbol{C} \otimes \frac{\partial}{\partial \theta_r} \boldsymbol{R}\right)\right) \\
\overset{tr(AB)=tr(BA)}{=} \quad & \operatorname{tr}\left((\boldsymbol{S_{\tilde{C}}} \otimes \boldsymbol{S_R} + \boldsymbol{I} \otimes \boldsymbol{I})^{-1} \left(\boldsymbol{U_\Sigma} \boldsymbol{S_\Sigma}^{\frac{1}{2}} \boldsymbol{U_{\tilde{C}}} \otimes \boldsymbol{U_R}\right)^{-1} \left(\boldsymbol{C} \otimes \frac{\partial}{\partial \theta_r} \boldsymbol{R}\right) \left(\boldsymbol{U_{\tilde{C}}}^\top \boldsymbol{S_\Sigma}^{\frac{1}{2}} \boldsymbol{U_\Sigma}^\top \otimes \boldsymbol{U_R}^\top\right)^{-1}\right) \\
\overset{(1.16)}{=} \quad & \operatorname{tr}\left((\boldsymbol{S_{\tilde{C}}} \otimes \boldsymbol{S_R} + \boldsymbol{I} \otimes \boldsymbol{I}) \left(\boldsymbol{U_{\tilde{C}}}^\top \boldsymbol{S_\Sigma}^{-\frac{1}{2}} \boldsymbol{U_\Sigma}^\top \boldsymbol{C} \boldsymbol{U_\Sigma} \boldsymbol{S_\Sigma}^{-\frac{1}{2}} \boldsymbol{U_{\tilde{C}}} \otimes \boldsymbol{U_R}^\top \left(\frac{\partial}{\partial \theta_r} \boldsymbol{R}\right) \boldsymbol{U_R}\right)\right) \\
\overset{\text{def}(\tilde{\boldsymbol{C}})}{=} \quad & \operatorname{tr}\left((\boldsymbol{S_{\tilde{C}}} \otimes \boldsymbol{S_R} + \boldsymbol{I} \otimes \boldsymbol{I}) \left(\boldsymbol{U_{\tilde{C}}}^\top \tilde{\boldsymbol{C}} \boldsymbol{U_{\tilde{C}}} \otimes \boldsymbol{U_R}^\top \left(\frac{\partial}{\partial \theta_r} \boldsymbol{R}\right) \boldsymbol{U_R}\right)\right) \\
\overset{tr(AB)=\operatorname{vec}(A)^\top \operatorname{vec}(B)}{=} \quad & \operatorname{diag}(\boldsymbol{S_{\tilde{C}}} \otimes \boldsymbol{S_R} + \boldsymbol{I} \otimes \boldsymbol{I})^\top \left[\operatorname{diag}(\boldsymbol{S_{\tilde{C}}}) \otimes \operatorname{diag}\left(\boldsymbol{U_R}^\top \left(\frac{\partial}{\partial \theta_r} \boldsymbol{R}\right) \boldsymbol{U_R}\right)\right]
\end{aligned} \tag{1.28}$$

The gradient of the squared form is given by:

$$\begin{aligned}
& \frac{\partial}{\partial \theta_r} \operatorname{vec}(\boldsymbol{Y}_r)^\top (\boldsymbol{C} \otimes \boldsymbol{R} + \boldsymbol{\Sigma} \otimes \boldsymbol{I})^{-1} \operatorname{vec}(\boldsymbol{Y}_r) \\
& = -\operatorname{vec}(\boldsymbol{Y}_r)^\top (\boldsymbol{C} \otimes \boldsymbol{R} + \boldsymbol{\Sigma} \otimes \boldsymbol{I})^{-1} \left(\boldsymbol{C} \otimes \frac{\partial}{\partial \theta_r} \boldsymbol{R}\right) (\boldsymbol{C} \otimes \boldsymbol{R} + \boldsymbol{\Sigma} \otimes \boldsymbol{I})^{-1} \operatorname{vec}(\boldsymbol{Y}_r) \\
& \overset{(1.27)}{=} -\operatorname{vec}\left(\tilde{\boldsymbol{Y}}_r\right)^\top \left(\boldsymbol{U_\Sigma} \boldsymbol{S_\Sigma}^{\frac{1}{2}} \boldsymbol{U_{\tilde{C}}} \otimes \boldsymbol{U_R}\right)^{-1} \left(\boldsymbol{C} \otimes \frac{\partial}{\partial \theta_r} \boldsymbol{R}\right) \left(\boldsymbol{U_{\tilde{C}}}^\top \boldsymbol{S_\Sigma}^{\frac{1}{2}} \boldsymbol{U_\Sigma}^\top \otimes \boldsymbol{U_R}^\top\right)^{-1} \operatorname{vec}\left(\tilde{\boldsymbol{Y}}_r\right) \\
& \overset{(1.16)}{=} -\operatorname{vec}\left(\tilde{\boldsymbol{Y}}_r\right) \left[\boldsymbol{U_{\tilde{C}}}^\top \boldsymbol{S_\Sigma}^{-\frac{1}{2}} \boldsymbol{U_\Sigma}^\top \boldsymbol{C} \boldsymbol{U_\Sigma} \boldsymbol{S_\Sigma}^{-\frac{1}{2}} \boldsymbol{U_{\tilde{C}}} \otimes \boldsymbol{U_R}^\top \left(\frac{\partial}{\partial \theta_r} \boldsymbol{R}\right) \boldsymbol{U_R}\right] \operatorname{vec}\left(\tilde{\boldsymbol{Y}}_r\right) \\
& \overset{\text{def}(\tilde{\boldsymbol{C}})}{=} -\operatorname{vec}\left(\tilde{\boldsymbol{Y}}_r\right)^\top \left[\boldsymbol{S_{\tilde{C}}} \otimes \boldsymbol{U_R}^\top \left(\frac{\partial}{\partial \theta_r} \boldsymbol{R}\right) \boldsymbol{U_R}\right] \operatorname{vec}\left(\tilde{\boldsymbol{Y}}_r\right) \\
& \overset{(1.17)}{=} -\operatorname{vec}\left(\tilde{\boldsymbol{Y}}_r\right)^\top \operatorname{vec}\left[\boldsymbol{U_R}^\top \left(\frac{\partial}{\partial \theta_r} \boldsymbol{R}\right) \boldsymbol{U_R} \tilde{\boldsymbol{Y}}_r \boldsymbol{S_{\tilde{C}}}\right]
\end{aligned} \tag{1.29}$$

##### 1.4.3 Estimation of the fixed effects

**Gradient Evaluation** We first evaluate the gradient with respect to a single fixed effect

$$
\begin{aligned}
& \frac{\partial}{[\partial \boldsymbol{B}_k]_{a,b}} \left( -\frac{1}{2} \mathrm{vec}(\boldsymbol{Y}_r)^\top (\boldsymbol{C} \otimes \boldsymbol{R} + \boldsymbol{\Sigma} \otimes \boldsymbol{I})^{-1} \mathrm{vec}(\boldsymbol{Y}_r) \right) \\
= \quad & -\mathrm{vec}(\boldsymbol{Y}_r)^\top (\boldsymbol{C} \otimes \boldsymbol{R} + \boldsymbol{\Sigma} \otimes \boldsymbol{I})^{-1} \mathrm{vec}\left( \frac{\partial \boldsymbol{Y}_r}{[\partial \boldsymbol{B}_k]_{a,b}} \right) \\
\overset{(1.27)}{=} \quad & -\mathrm{vec}\left( \tilde{\boldsymbol{Y}}_r \right)^\top \left( \boldsymbol{U}_{\tilde{\boldsymbol{C}}}^\top \boldsymbol{S}_{\boldsymbol{\Sigma}}^{-\frac{1}{2}} \boldsymbol{U}_{\boldsymbol{\Sigma}}^\top \otimes \boldsymbol{U}_{\boldsymbol{R}}^\top \right) \mathrm{vec}\left( \frac{\partial \boldsymbol{Y}_r}{[\partial \boldsymbol{B}_k]_{a,b}} \right) \\
\overset{(1.17)}{=} \quad & -\mathrm{vec}\left( \tilde{\boldsymbol{Y}}_r \right) \mathrm{vec}\left( \boldsymbol{U}_{\boldsymbol{R}}^\top \left( \frac{\partial \boldsymbol{Y}_r}{[\partial \boldsymbol{B}_k]_{a,b}} \right) \boldsymbol{U}_{\boldsymbol{\Sigma}} \boldsymbol{S}_{\boldsymbol{\Sigma}}^{-\frac{1}{2}} \boldsymbol{U}_{\tilde{\boldsymbol{C}}} \right) \\
= \quad & -\mathrm{vec}\left( \tilde{\boldsymbol{Y}}_r \right)^\top \mathrm{vec}\left( \boldsymbol{U}_{\boldsymbol{R}}^\top \left( [\boldsymbol{X}_k]_{:,a} [\boldsymbol{A}_k]_{:,b}^\top \right) \boldsymbol{U}_{\boldsymbol{\Sigma}} \boldsymbol{S}_{\boldsymbol{\Sigma}}^{-\frac{1}{2}} \boldsymbol{U}_{\tilde{\boldsymbol{C}}} \right) \\
\overset{tr(AB)=\mathrm{vec}(A)^\top \mathrm{vec}(B)}{=} \quad & -\mathrm{tr}\left( \tilde{\boldsymbol{Y}}_r^\top \boldsymbol{U}_{\boldsymbol{R}}^\top [\boldsymbol{X}_k]_{:,a} [\boldsymbol{A}_k]_{:,b}^\top \boldsymbol{U}_{\boldsymbol{\Sigma}} \boldsymbol{S}_{\boldsymbol{\Sigma}}^{-\frac{1}{2}} \boldsymbol{U}_{\tilde{\boldsymbol{C}}} \right) \\
\overset{tr(AB)=tr(A)tr(B)}{=} \quad & -\mathrm{tr}\left( [\boldsymbol{A}_k]_{:,b}^\top \boldsymbol{U}_{\boldsymbol{\Sigma}} \boldsymbol{S}_{\boldsymbol{\Sigma}}^{-\frac{1}{2}} \boldsymbol{U}_{\tilde{\boldsymbol{C}}} \tilde{\boldsymbol{Y}}_r^\top \boldsymbol{U}_{\boldsymbol{R}}^\top [\boldsymbol{X}_k]_{:,a} \right) \\
\overset{tr(A)=tr(A^\top)}{=} \quad & -[\boldsymbol{X}_k]_{:,a}^\top \boldsymbol{U}_{\boldsymbol{R}} \tilde{\boldsymbol{Y}}_r \boldsymbol{U}_{\tilde{\boldsymbol{C}}}^\top \boldsymbol{S}_{\boldsymbol{\Sigma}}^{-\frac{1}{2}} \boldsymbol{U}_{\boldsymbol{\Sigma}}^\top [\boldsymbol{A}_k]_{:,b} \qquad (1.30)
\end{aligned}
$$

When stacking together the derivatives for all $a$ and $b$, the gradient with respect to all entries of $\boldsymbol{B}_k$ follows as

$$
\begin{aligned}
& \frac{\Delta}{\Delta \boldsymbol{B}_k} \left( -\frac{1}{2} \mathrm{vec}(\boldsymbol{Y}_r)^\top (\boldsymbol{C} \otimes \boldsymbol{R} + \boldsymbol{\Sigma} \otimes \boldsymbol{I})^{-1} \mathrm{vec}(\boldsymbol{Y}_r) \right) \\
&= - \boldsymbol{X}_k^\top \boldsymbol{U}_{\boldsymbol{R}} \tilde{\boldsymbol{Y}}_r \boldsymbol{U}_{\tilde{\boldsymbol{C}}}^\top \boldsymbol{S}_{\boldsymbol{\Sigma}}^{-\frac{1}{2}} \boldsymbol{U}_{\boldsymbol{\Sigma}}^\top \boldsymbol{A}_k \qquad (1.31)
\end{aligned}
$$

**Closed form maximum likelihood estimates** First, we rewrite the mean function by concatenating the Kronecker products of the design matrix

$$\Phi = [\boldsymbol{A}_1 \otimes \boldsymbol{X}_1, \ldots, \boldsymbol{A}_J \otimes \boldsymbol{X}_J] \qquad (1.32)$$

and concatenating the fixed effects

$$\boldsymbol{\beta} = \begin{pmatrix} \mathrm{vec}(\boldsymbol{B}_1) \\ \ldots \\ \mathrm{vec}(\boldsymbol{B}_J) \end{pmatrix}, \qquad (1.33)$$

where $\boldsymbol{\Phi}$ is a $NP \times DM$ matrix and $\boldsymbol{\beta}$ is a $DM \times 1$ vector, with $D = \sum_j D_j$ and $M = \sum_j M_j$. It is easy to show that $\sum_{j=1}^J (\boldsymbol{A}_j \otimes \boldsymbol{X}_j) \mathrm{vec}(\boldsymbol{B}_j) = \boldsymbol{\Phi}\boldsymbol{\beta}$.

By setting the gradient of the log-likelihood with respect to the weight vector to zero, we can then obtain a closed form solution of the maximum-likelihood estimate $\boldsymbol{\beta}_M$:

$$\frac{\nabla}{\nabla \boldsymbol{\beta}_{\mathrm{M}}} \left( -\frac{1}{2} (\mathrm{vec}(\boldsymbol{Y}) - \Phi\boldsymbol{\beta})^\top (\boldsymbol{C} \otimes \boldsymbol{R} + \boldsymbol{\Sigma} \otimes \boldsymbol{I})^{-1} (\mathrm{vec}(\boldsymbol{Y}) - \Phi\boldsymbol{\beta}) \right) = \boldsymbol{0} \qquad (1.34)$$

$$\Phi^\top (\boldsymbol{C} \otimes \boldsymbol{R} + \boldsymbol{\Sigma} \otimes \boldsymbol{I})^{-1} \Phi\boldsymbol{\beta}_{\mathrm{M}} - \Phi^\top (\boldsymbol{C} \otimes \boldsymbol{R} + \boldsymbol{\Sigma} \otimes \boldsymbol{I})^{-1} \mathrm{vec}(\boldsymbol{Y}) = \boldsymbol{0} \qquad (1.35)$$

$$\Phi^\top (\boldsymbol{C} \otimes \boldsymbol{R} + \boldsymbol{\Sigma} \otimes \boldsymbol{I})^{-1} \Phi\boldsymbol{\beta} = \Phi^\top (\boldsymbol{C} \otimes \boldsymbol{R} + \boldsymbol{\Sigma} \otimes \boldsymbol{I})^{-1} \mathrm{vec}(\boldsymbol{Y}) \qquad (1.36)$$

$$\boldsymbol{\beta}_{\mathrm{M}} = \Big(\underbrace{\Phi^{\top}\left(\boldsymbol{C}\otimes\boldsymbol{R}+\boldsymbol{\Sigma}\otimes\boldsymbol{I}\right)^{-1}\Phi}_{\text{left term}}\Big)^{-1}\underbrace{\Phi^{\top}\left(\boldsymbol{C}\otimes\boldsymbol{R}+\ \boldsymbol{\Sigma}\otimes\boldsymbol{I}\right)^{-1}\operatorname{vec}\left(\boldsymbol{Y}\right)}_{\text{right term}} \tag{1.37}$$

We first compute the right term with respect to the $j^{\text{th}}$ block.

$$\begin{aligned}
&\left(\boldsymbol{A}_j\otimes\boldsymbol{X}_j\right)^{\top}\left(\boldsymbol{C}\otimes\boldsymbol{R}+\boldsymbol{\Sigma}\otimes\boldsymbol{I}\right)^{-1}\operatorname{vec}\left(\boldsymbol{Y}\right)\\
\overset{(1.27)}{=}&\left(\boldsymbol{A}_j\otimes\boldsymbol{X}_j\right)^{\top}\left(\boldsymbol{U}_{\boldsymbol{\Sigma}}\boldsymbol{S}_{\boldsymbol{\Sigma}}{}^{-\frac{1}{2}}\boldsymbol{U}_{\tilde{\boldsymbol{C}}}\otimes\boldsymbol{U}_{\boldsymbol{R}}\right)\operatorname{vec}\left(\boldsymbol{D}\odot\boldsymbol{U}_{\boldsymbol{R}}^{\top}\boldsymbol{Y}_r\boldsymbol{U}_{\boldsymbol{\Sigma}}\boldsymbol{S}_{\boldsymbol{\Sigma}}{}^{-\frac{1}{2}}\boldsymbol{U}_{\tilde{\boldsymbol{C}}}\right)\\
\overset{(1.16)}{=}&\left(\boldsymbol{A}_j^{\top}\boldsymbol{U}_{\boldsymbol{\Sigma}}\boldsymbol{S}_{\boldsymbol{\Sigma}}{}^{-\frac{1}{2}}\boldsymbol{U}_{\tilde{\boldsymbol{C}}}\otimes\boldsymbol{X}_j^{\top}\boldsymbol{U}_{\boldsymbol{R}}\right)\operatorname{vec}\left(\boldsymbol{D}\odot\boldsymbol{U}_{\boldsymbol{R}}^{\top}\boldsymbol{Y}_r\boldsymbol{U}_{\boldsymbol{\Sigma}}\boldsymbol{S}_{\boldsymbol{\Sigma}}{}^{-\frac{1}{2}}\boldsymbol{U}_{\tilde{\boldsymbol{C}}}\right)\\
\overset{(1.17)}{=}&\operatorname{vec}\left[\boldsymbol{X}_j^{\top}\boldsymbol{U}_{\boldsymbol{R}}\left(\boldsymbol{D}\odot\boldsymbol{U}_{\boldsymbol{R}}^{\top}\boldsymbol{Y}_r\boldsymbol{U}_{\boldsymbol{\Sigma}}\boldsymbol{S}_{\boldsymbol{\Sigma}}{}^{-\frac{1}{2}}\boldsymbol{U}_{\tilde{\boldsymbol{C}}}\right)\boldsymbol{U}_{\tilde{\boldsymbol{C}}}^{\top}\boldsymbol{S}_{\boldsymbol{\Sigma}}{}^{-\frac{1}{2}}\boldsymbol{U}_{\boldsymbol{\Sigma}}^{\top}\boldsymbol{A}_j\right]
\end{aligned} \tag{1.38}$$

By concatenating the blocks, we obtain

$$\Phi^{\top}\left(\boldsymbol{C}\otimes\boldsymbol{R}+\sigma^2\boldsymbol{I}\right)^{-1}\operatorname{vec}\left(\boldsymbol{Y}\right)=\begin{bmatrix}\operatorname{vec}\left(\boldsymbol{X}_1^{T}\boldsymbol{U}_{\boldsymbol{R}}\left(\boldsymbol{D}\odot\boldsymbol{U}_{\boldsymbol{R}}^{\top}\boldsymbol{Y}_r\boldsymbol{U}_{\boldsymbol{\Sigma}}\boldsymbol{S}_{\boldsymbol{\Sigma}}{}^{-\frac{1}{2}}\boldsymbol{U}_{\tilde{\boldsymbol{C}}}\right)\boldsymbol{U}_{\tilde{\boldsymbol{C}}}^{\top}\boldsymbol{S}_{\boldsymbol{\Sigma}}{}^{-\frac{1}{2}}\boldsymbol{U}_{\boldsymbol{\Sigma}}^{\top}\boldsymbol{A}_1\right)\\ \vdots\\ \operatorname{vec}\left(\boldsymbol{X}_J^{T}\boldsymbol{U}_{\boldsymbol{R}}\left(\boldsymbol{D}\odot\boldsymbol{U}_{\boldsymbol{R}}^{\top}\boldsymbol{Y}_r\boldsymbol{U}_{\boldsymbol{\Sigma}}\boldsymbol{S}_{\boldsymbol{\Sigma}}{}^{-\frac{1}{2}}\boldsymbol{U}_{\tilde{\boldsymbol{C}}}\right)\boldsymbol{U}_{\tilde{\boldsymbol{C}}}^{\top}\boldsymbol{S}_{\boldsymbol{\Sigma}}{}^{-\frac{1}{2}}\boldsymbol{U}_{\boldsymbol{\Sigma}}^{\top}\boldsymbol{A}_J\right)\end{bmatrix} \tag{1.39}$$

For the left term, we need to invert the block matrix $\Phi^{\top}\left(\boldsymbol{C}\otimes\boldsymbol{R}+\boldsymbol{\Sigma}\otimes\boldsymbol{I}\right)^{-1}\Phi\in\mathbb{R}^{DM\times DM}$. We start with computing the $(i,j)^{\text{th}}$ block involving the $i^{\text{th}}$ and the $j^{\text{th}}$ fixed effects:

$$\begin{aligned}
&\left(\boldsymbol{A}_i\otimes\boldsymbol{X}_i\right)^{\top}\left(\boldsymbol{C}\otimes\boldsymbol{R}+\boldsymbol{\Sigma}\otimes\boldsymbol{I}\right)^{-1}\left(\boldsymbol{A}_j\otimes\boldsymbol{X}_j\right)\\
\overset{(1.22)}{=}\;&\left(\boldsymbol{A}_j\otimes\boldsymbol{X}_j\right)^{\top}\left[\left(\boldsymbol{U}_{\boldsymbol{\Sigma}}\boldsymbol{S}_{\boldsymbol{\Sigma}}^{\frac{1}{2}}\boldsymbol{U}_{\tilde{\boldsymbol{C}}}\otimes\boldsymbol{U}_{\boldsymbol{R}}\right)\left(\boldsymbol{S}_{\tilde{\boldsymbol{C}}}\otimes\boldsymbol{S}_{\boldsymbol{R}}+\boldsymbol{I}\otimes\boldsymbol{I}\right)\left(\boldsymbol{U}_{\tilde{\boldsymbol{C}}}^{\top}\boldsymbol{S}_{\boldsymbol{\Sigma}}^{\frac{1}{2}}\boldsymbol{U}_{\boldsymbol{\Sigma}}{}^{\top}\otimes\boldsymbol{U}_{\boldsymbol{R}}^{\top}\right)\right]^{-1}\left(\boldsymbol{A}_j\otimes\boldsymbol{X}_j\right)\\
=\;&\left(\boldsymbol{A}_i\otimes\boldsymbol{X}_i\right)^{\top}\left(\boldsymbol{U}_{\boldsymbol{\Sigma}}\boldsymbol{S}_{\boldsymbol{\Sigma}}{}^{-\frac{1}{2}}\boldsymbol{U}_{\tilde{\boldsymbol{C}}}\otimes\boldsymbol{U}_{\boldsymbol{R}}\right)\left(\boldsymbol{S}_{\tilde{\boldsymbol{C}}}\otimes\boldsymbol{S}_{\boldsymbol{R}}+\boldsymbol{I}\otimes\boldsymbol{I}\right)^{-1}\left(\boldsymbol{U}_{\tilde{\boldsymbol{C}}}^{\top}\boldsymbol{S}_{\boldsymbol{\Sigma}}{}^{-\frac{1}{2}}\boldsymbol{U}_{\boldsymbol{\Sigma}}{}^{\top}\otimes\boldsymbol{U}_{\boldsymbol{R}}^{\top}\right)\left(\boldsymbol{A}_j\otimes\boldsymbol{X}_j\right)\\
\overset{(1.16)}{=}\;&\left(\boldsymbol{A}_i^{\top}\boldsymbol{U}_{\boldsymbol{\Sigma}}\boldsymbol{S}_{\boldsymbol{\Sigma}}{}^{-\frac{1}{2}}\boldsymbol{U}_{\tilde{\boldsymbol{C}}}\otimes\boldsymbol{X}_i^{\top}\boldsymbol{U}_{\boldsymbol{R}}\right)\left(\boldsymbol{S}_{\tilde{\boldsymbol{C}}}\otimes\boldsymbol{S}_{\boldsymbol{R}}+\boldsymbol{I}\otimes\boldsymbol{I}\right)^{-1}\left(\boldsymbol{U}_{\tilde{\boldsymbol{C}}}^{\top}\boldsymbol{S}_{\boldsymbol{\Sigma}}{}^{-\frac{1}{2}}\boldsymbol{U}_{\boldsymbol{\Sigma}}{}^{\top}\boldsymbol{A}_j\otimes\boldsymbol{U}_{\boldsymbol{R}}^{\top}\boldsymbol{X}_j\right)\\
=\;&\sum_{c=1}^{P}\left(\left[\boldsymbol{A}_i^{\top}\boldsymbol{U}_{\boldsymbol{\Sigma}}\boldsymbol{S}_{\boldsymbol{\Sigma}}{}^{-\frac{1}{2}}\boldsymbol{U}_{\tilde{\boldsymbol{C}}}\right]_{:,c}\otimes\boldsymbol{X}_i^{\top}\boldsymbol{U}_{\boldsymbol{R}}\right)\left(\boldsymbol{S}_{\tilde{\boldsymbol{C}}}[c,c]\boldsymbol{S}_{\boldsymbol{R}}+\boldsymbol{I}\right)^{-1}\left(\left[\boldsymbol{U}_{\tilde{\boldsymbol{C}}}^{\top}\boldsymbol{S}_{\boldsymbol{\Sigma}}{}^{-\frac{1}{2}}\boldsymbol{U}_{\boldsymbol{\Sigma}}{}^{\top}\boldsymbol{A}_j\right]_{c,:}\otimes\boldsymbol{U}_{\boldsymbol{R}}^{\top}\boldsymbol{X}_j\right)\\
\overset{(1.16)}{=}\;&\sum_{c=1}^{P}\left(\left[\boldsymbol{A}_i^{\top}\boldsymbol{U}_{\boldsymbol{\Sigma}}\boldsymbol{S}_{\boldsymbol{\Sigma}}{}^{-\frac{1}{2}}\boldsymbol{U}_{\tilde{\boldsymbol{C}}}\right]_{:,c}\otimes\boldsymbol{X}_i^{\top}\boldsymbol{U}_{\boldsymbol{R}}\right)\left(\left[\boldsymbol{U}_{\tilde{\boldsymbol{C}}}^{\top}\boldsymbol{S}_{\boldsymbol{\Sigma}}{}^{-\frac{1}{2}}\boldsymbol{U}_{\boldsymbol{\Sigma}}{}^{\top}\boldsymbol{A}_j\right]_{c,:}\otimes\left(\boldsymbol{S}_{\tilde{\boldsymbol{C}}}[c,c]\boldsymbol{S}_{\boldsymbol{R}}+\boldsymbol{I}\right)^{-1}\boldsymbol{U}_{\boldsymbol{R}}^{\top}\boldsymbol{X}_j\right)\\
\overset{(1.16)}{=}\;&\sum_{c=1}^{P}\left(\left[\boldsymbol{A}_i^{\top}\boldsymbol{U}_{\boldsymbol{\Sigma}}\boldsymbol{S}_{\boldsymbol{\Sigma}}{}^{-\frac{1}{2}}\boldsymbol{U}_{\tilde{\boldsymbol{C}}}\right]_{:,c}\left[\boldsymbol{U}_{\tilde{\boldsymbol{C}}}^{\top}\boldsymbol{S}_{\boldsymbol{\Sigma}}{}^{-\frac{1}{2}}\boldsymbol{U}_{\boldsymbol{\Sigma}}{}^{\top}\boldsymbol{A}_j\right]_{c,:}\otimes\boldsymbol{X}_i^{\top}\boldsymbol{U}_{\boldsymbol{R}}\left(\boldsymbol{S}_{\tilde{\boldsymbol{C}}}[c,c]\boldsymbol{S}_{\boldsymbol{R}}+\boldsymbol{I}\right)^{-1}\boldsymbol{U}_{\boldsymbol{R}}^{\top}\boldsymbol{X}_j\right)
\end{aligned} \tag{1.40}$$

Evaluating the term takes $O(PM^2+PD^2N)$ time. If the number of samples is smaller than the number of traits ($N<P$), we can also explicitly sum over the samples leading to a runtime of $O(ND^2+NM^2P)$. Inverting the left term has a runtime complexity of $O(D^3M^3)$.

##### 1.4.4 Predictions

The mean predictor can be evaluated efficiently as follows

$$
\begin{aligned}
\operatorname{vec}(\boldsymbol{M}^{*}) &= \sum_{j=1}^{J} \left(\boldsymbol{A}_j \otimes \boldsymbol{X}_j^{\star}\right) \operatorname{vec}(\boldsymbol{B}_j) + \left(\boldsymbol{C} \otimes \boldsymbol{R}^{\star}\right) \left(\boldsymbol{C} \otimes \boldsymbol{R} + \boldsymbol{\Sigma} \otimes \boldsymbol{I}\right)^{-1} \operatorname{vec}\left(\boldsymbol{Y} - \sum_{j=1}^{J} \left(\boldsymbol{A}_j \otimes \boldsymbol{X}_j\right) \operatorname{vec}(\boldsymbol{B}_j)\right) \\
&\overset{\operatorname{def}(Y_r)}{=} \sum_{j=1}^{J} \boldsymbol{X}_j^{\star} \boldsymbol{B}_j \boldsymbol{A}_j^{T} + \left(\boldsymbol{C} \otimes \boldsymbol{R}^{\star}\right) \left(\boldsymbol{C} \otimes \boldsymbol{R} + \boldsymbol{\Sigma} \otimes \boldsymbol{I}\right)^{-1} \operatorname{vec}(\boldsymbol{Y}_r) \\
&\overset{(1.27)}{=} \sum_{j=1}^{J} \boldsymbol{X}_j^{\star} \boldsymbol{B}_j \boldsymbol{A}_j^{T} + \left(\boldsymbol{C} \otimes \boldsymbol{R}^{\star}\right) \left(\boldsymbol{U}_{\boldsymbol{\Sigma}} \boldsymbol{S}_{\boldsymbol{\Sigma}}{}^{-\frac{1}{2}} \boldsymbol{U}_{\tilde{\boldsymbol{C}}} \otimes \boldsymbol{U}_{\boldsymbol{R}}\right) \operatorname{vec}\left(\tilde{\boldsymbol{Y}}_r\right) \\
&\overset{(1.16)}{=} \sum_{j=1}^{J} \boldsymbol{X}_j^{\star} \boldsymbol{B}_j \boldsymbol{A}_j^{T} + \left(\boldsymbol{C} \boldsymbol{U}_{\boldsymbol{\Sigma}} \boldsymbol{S}_{\boldsymbol{\Sigma}}{}^{-\frac{1}{2}} \boldsymbol{U}_{\tilde{\boldsymbol{C}}} \otimes \boldsymbol{R}^{\star} \boldsymbol{U}_{\boldsymbol{R}}\right) \operatorname{vec}\left(\tilde{\boldsymbol{Y}}_r\right) \\
&\overset{(1.17)}{=} \sum_{j=1}^{J} \boldsymbol{X}_j^{\star} \boldsymbol{B}_j \boldsymbol{A}_j^{T} + \operatorname{vec}\left[\boldsymbol{R}^{\star} \boldsymbol{U}_{\boldsymbol{R}} \tilde{\boldsymbol{Y}}_r \left(\boldsymbol{C} \boldsymbol{U}_{\boldsymbol{\Sigma}} \boldsymbol{S}_{\boldsymbol{\Sigma}}{}^{-\frac{1}{2}} \boldsymbol{U}_{\tilde{\boldsymbol{C}}}\right)^{\top}\right] \\
&= \sum_{j=1}^{J} \boldsymbol{X}_j^{\star} \boldsymbol{B}_j \boldsymbol{A}_j^{T} + \operatorname{vec}\left(\boldsymbol{R}^{\star} \boldsymbol{U}_{\boldsymbol{R}} \tilde{\boldsymbol{Y}}_r \boldsymbol{U}_{\tilde{\boldsymbol{C}}}^{\top} \boldsymbol{S}_{\boldsymbol{\Sigma}}{}^{-\frac{1}{2}} \boldsymbol{U}_{\boldsymbol{\Sigma}}^{\top} \boldsymbol{C}^{\top}\right) \qquad (1.41)
\end{aligned}
$$

#### 1.5 Implementation details

###### OOP design

###### code example

### 2 Multivariate linear mixed models for statistical genetics

#### 2.1 Introduction

In the most general formulation LIMIX models an $N \times P$ matrix $\mathbf{Y}$ of $N$ samples for each of $P$ phenotypes by a multivariate linear mixed model with $J$ fixed effects $\{\boldsymbol{F}_j\}$ and $I$ random effects $\{\boldsymbol{U}_i\}$

$$\boldsymbol{Y} = \sum_{j=1}^{J} \boldsymbol{F}_j + \sum_{i=1}^{I} \boldsymbol{U}_i. \tag{2.1}$$

Each of the fixed effects and random effects is Kronecker structured. In particular, any fixed effect $\boldsymbol{F}_i$ can be written as $(\boldsymbol{A}_i \otimes \boldsymbol{X}_i) \operatorname{vec}(\boldsymbol{B}_i)$ where $\boldsymbol{A}_i \in \mathbb{R}^{P \times M}$ is the design matrix of the phenotypic effects, $\boldsymbol{X}_i \in \mathbb{R}^{N \times D}$ is the design matrix of the sample-specific effects, and $\boldsymbol{B}_i \in \mathbb{R}^{M \times D}$ is the matrix of the effect sizes; while any random effect $\boldsymbol{U}_i$ is matrix-variate normal distributed, $\boldsymbol{U}_i \sim \mathcal{N}_{NM}(\mathbf{0}\,;\, \boldsymbol{R}_i\,,\, \boldsymbol{C}_i)$, with row covariance matrix $\boldsymbol{R}_i \in \mathbb{R}^{N \times N}$ and column covariance matrix $\boldsymbol{C}_i \in \mathbb{R}^{P \times P}$. $\boldsymbol{R}_i$ and $\boldsymbol{C}_i$ can be interpreted as sample-to-sample and trait-to-trait relatedness matrices due to contribution $i$ respectively.

All the models described by equation (2.1) can be dynamically built using LIMIX and subsequently fitted to the data using the optimization framework we introduced in Section 1. In the following subsection we show how some commonly used mixed models for genetic studies can be written as in (2.1). Afterwards we review common types of genetic studies that are supported by LIMIX. In Section 2.2 we present the framework made available by LIMIX to dissect the phenotypic variability across different sources and make out-of-sample phenotype predictions. As discussed in the main text, generalization performance measured by out-of-sample prediction can be used as a model selection criterion, given the wealth of models made available by LIMIX. In Section 2.3 we describe models for genome-wide association studies (GWAS) and show how they can be extended to build a multi-locus multivariate model. Finally, in Section 2.4 we discuss the PANAMA module in LIMIX, which can be used to infer non-observed (hidden) covariates so that they can be accounted for in GWAS and variance decomposition.

##### 2.1.1 Some recently proposed MTMM

A particular, yet multivariate case of this general model is when the same design is used for all fixed effects and only two random effects describing the genetic and non-genetic contributions to the phenotype are considered:

$$\boldsymbol{Y} = (\boldsymbol{A} \otimes \boldsymbol{X}) \operatorname{vec}(\boldsymbol{B}) + \boldsymbol{U} + \boldsymbol{\Psi}, \quad \boldsymbol{U} \sim \mathcal{N}_{NM}(\mathbf{0}\,;\, \boldsymbol{R}\,,\, \boldsymbol{C}), \; \boldsymbol{\Psi} \sim \mathcal{N}_{NM}(\mathbf{0}\,;\, \boldsymbol{I}\,,\, \Sigma) \tag{2.2}$$

where $\boldsymbol{R}$ is the kinship matrix, which describes the genetic relatedness between the samples, $\boldsymbol{C}$ describes the genetic relatedness between the phenotypes, and $\boldsymbol{\Sigma}$ the relatedness between the phenotypes due to shared non-gentic effects, i.e. noise. Similar models have been recently employed in genetic analysis of multiple phenotypes [Zhou and Stephens, 2014, Korte et al., 2012, Rakitsch et al., Stegle et al., 2011b].

Considering the univariate version of the model in (2.2), the trait design matrix in the fixed effect collapses to 1 while the trait-to-trait covariance matrices reduce to single variance components. Indicating with $\boldsymbol{y}$, $\boldsymbol{\beta}$, $\boldsymbol{u}$, $\boldsymbol{\psi}$, $\sigma_g^2$ and $\sigma_n^2$ the univariate versions of $\boldsymbol{Y}$, $\boldsymbol{B}$, $\boldsymbol{U}$, $\boldsymbol{\Psi}$, $\boldsymbol{C}^2$ and $\boldsymbol{\Sigma}^2$ the model reduces to

$$\boldsymbol{y} = \boldsymbol{X}\boldsymbol{\beta} + \boldsymbol{u} + \boldsymbol{\psi}, \quad \boldsymbol{u} \sim \mathcal{N}\left(\mathbf{0},\, \sigma_g^2 \boldsymbol{R}\right), \; \boldsymbol{\Psi} \sim \mathcal{N}\left(\mathbf{0},\, \sigma_e^2 \boldsymbol{I}\right) \tag{2.3}$$

Very similar models have been widely used in genetic studies for heritability estimation and GWAS [Kang et al., 2010, Lippert et al., 2011, Listgarten et al., 2013, Segura et al., 2012].

Instead of considering the effect from single variants as in (2.3), one can consider the effects from all SNPs in a region (e.g., a gene or a chromosome). This effect, $\boldsymbol{r}$, can be modelled as random effect whose covariance matrix $\boldsymbol{S}$ is a region-based genetic relatedness matrix:

$$\boldsymbol{y} = \boldsymbol{r} + \boldsymbol{u} + \boldsymbol{\psi}, \quad \boldsymbol{r} \sim \mathcal{N}\left(\boldsymbol{0},\, \sigma_r^2 \boldsymbol{S}\right), \boldsymbol{u} \sim \mathcal{N}\left(\boldsymbol{0},\, \sigma_g^2 \boldsymbol{R}\right),\ \boldsymbol{\Psi} \sim \mathcal{N}\left(\boldsymbol{0},\, \sigma_e^2 \boldsymbol{I}\right) \tag{2.4}$$

Similar ideas and models have been employed for general set tests [Listgarten et al., 2013], rare variant testing [Wu et al., 2011], and heritability estimation [Yang et al., 2010].

#### 2.2 Variance Decomposition

In this section, we describe the variance decomposition framework made available by LIMIX that allows for dissecting the phenotypic variability across different sources for univariate (subsection 2.2.1) and multivariate analysis (subsection 2.2.2). In the same framework, LIMIX makes also available a tool for out-of-sample predictions, which we discuss in subsection 2.2.3.

##### 2.2.1 Single-trait variance decomposition

In the model for univariate variance decomposition, fixed effects reflects reflect covariates while random effects contributions from different sets of variants (e.g., different chromosomes or local and distal genetic contributions). The covariance matrices of these contributions are empirical covariance matrices describing genetic relatedness based on the different sets. The approach can be generalized to include non-genetic random effects whose covariance matrices are known *a priori*, allowing for correction of covariates and joint estimation of genetic and non-genetic contributions to the phenotypic variability.

The model can be written as:

$$\boldsymbol{y} = \boldsymbol{X}\boldsymbol{\beta} + \sum_i \boldsymbol{u}_i + \boldsymbol{\psi}, \quad \boldsymbol{u}_i \sim \mathcal{N}\left(\boldsymbol{0},\, \sigma_i^2 \boldsymbol{R}_i\right),\ \boldsymbol{\Psi} \sim \mathcal{N}\left(\boldsymbol{0},\, \sigma_e^2 \boldsymbol{I}\right) \tag{2.5}$$

where $\boldsymbol{R}_i$ is the covariance matrix for contribution $i$. If $\boldsymbol{R}$ is a genetic contribution from set of SNPs $\mathcal{S}$, with a bit of abuse of notation we can write

$$\boldsymbol{R} = \frac{1}{C} \boldsymbol{W}_{:,\mathcal{S}} \boldsymbol{W}_{:,\mathcal{S}}{}^T \tag{2.6}$$

where $C = \frac{1}{N}\mathrm{tr}\left(\boldsymbol{W}_{:,\mathcal{S}_i} \boldsymbol{W}_{:,\mathcal{S}_i}{}^T\right)$. The parameters of the model are the fixed effects $\boldsymbol{\beta}$ and the variance components $\sigma_i^2$ and $\sigma_e^2$.

Variance explained by the fixed effect $d$ can be retrieved by considering $\mathrm{var}\left(\boldsymbol{X}_{d,:}\boldsymbol{\beta}\right)$. P: think the following is superfluous Since one is usually interested in the fraction of total variance explained by each contribution rather than absolute variance contribution, variance components are normalized to sum up to 1.

##### 2.2.2 Multiple-trait variance decomposition

Using the notation introduced in Section 2.1, the general model for multivariate variance decomposition can be written as

$$\boldsymbol{Y} = \sum_i \left(\boldsymbol{A}_i \otimes \boldsymbol{X}_i\right) \mathrm{vec}\left(\boldsymbol{B}_i\right) + \sum_i \boldsymbol{U}_i + \boldsymbol{\Psi}, \qquad \boldsymbol{U}_i \sim \mathcal{N}_{NM}\left(\boldsymbol{0}\,;\, \boldsymbol{R}_i\,,\, \boldsymbol{C}_i(\boldsymbol{\alpha}_i)\right),\ \boldsymbol{\Psi} \sim \mathcal{N}_{NM}\left(\boldsymbol{0}\,;\, \boldsymbol{I}\,,\, \boldsymbol{\Sigma}\left(\boldsymbol{\alpha}_{\boldsymbol{\Sigma}}\right)\right) \tag{2.7}$$

where $\boldsymbol{U}_i$ indicates the genetic effect from random effect $i$ with trait-to-trait covariance matrix $\boldsymbol{C}_i$ and sample-to-sample covariance matrix $\boldsymbol{R}_i$ introduced in (2.6), $\boldsymbol{\Psi}$ denotes the non-genetic contribution. We have indicated with $\boldsymbol{\alpha}_i$, $\boldsymbol{\alpha}_{\boldsymbol{\Sigma}}$ the parameters of the trait-to-trait covariance matrices, which are estimated from the data. The parameters of the model are again the weight matrix of fixed effects $\boldsymbol{B}$ and the variance components $\boldsymbol{\alpha} = \{\boldsymbol{\alpha}_1, ..., \boldsymbol{\alpha}_{\boldsymbol{\Sigma}}\}$. Although LIMIX handles incomplete designs and cases where more than 2 random effect terms are involved, in the case of 2 random effect terms and without missing values in the phenotype matrix $\mathbf{Y}$ LIMIX exploits the Kronecker structure in the model and implements the mathematical tricks discussed in the previous chapter to conduct fast estimation of parameters.

In the most general model the trait-to-trait covariance matrices are general semi-definite positive matrices (*freeform* matrices) parametrized by $\frac{1}{2}P(P+1)$ real values. However, such a general form may lead to overfitting as the number of parameters increases quadratically in the number of traits while the data only linearly. LIMIX circumvents this problem in two ways:

- it makes available to the user a set of different covariance matrices to perform optimization of the $P \times P$ covariance matrices;
- it allows the user to introduce a regularization on the covariance parameters

For a technical discussion about the different covariance matrices and possible regularization schemes made available by LIMIX we refer the user to the previous chapter, while here we discuss utility and interpretation of these. First of all, we highlight that using a diagonal covariance matrix as trait-to-trait covariance matrix for random effect $i$ is equivalent to assume that term $i$ does not contribute to trait-to-trait correlations. A sum of a rank $R$ matrix and spherical residual $\boldsymbol{C}(\boldsymbol{\theta}) = \boldsymbol{A}\boldsymbol{A}^T + \sigma^2 \boldsymbol{I}$ (*lowrank-id*), where $\boldsymbol{A} \in \mathbb{R}^{P,R}$ and $\boldsymbol{\theta} = \left\{\text{vec}\left(\boldsymbol{A}\right), \sigma^2\right\}$, might help address overfitting problems as the number of parameters scales linearly with the number of traits. On the other hand, this parametrisation offers limited flexibility for the diagonal elements especially for low $R$, yielding biased heritability estimates. A more flexible form is the sum of a low-rank term and a trait-specific independent contribution $\boldsymbol{C}(\boldsymbol{\theta}) = \boldsymbol{A}\boldsymbol{A}^T + \text{diag}(\boldsymbol{c})$ (*lowrank-diag*), where $\boldsymbol{\theta} = \{\text{vec}\left(\boldsymbol{A}\right), \boldsymbol{c}\}$ and $\boldsymbol{c} \in \mathbb{R}^P$. The penalization on the off-diagonal entries of the trait-trait covariance matrix, or its inverse, bridges a fully independent model (*diagonal* matrix) to the unpenalised model. To select the extent of the penalization, one can use cross validation exploiting the phenotype prediction tools discussed in next subsection (2.2.3).

To dissect the contribution of each random effect term into shared and trait-specific effects, we decompose the trait-to-trait covariance matrix $\boldsymbol{C}_i$ into the biggest semi-positive definite block matrix contained in $\boldsymbol{C}_i$ and a residual: $\boldsymbol{C} = a^2 \mathbf{1}_{PP} + \boldsymbol{R}$, where $\mathbf{1}_{PP}$ denotes a $P \times P$ matrix of ones. Given this representation, we can then interpret $a^2$ and $\frac{1}{P}\text{tr}\boldsymbol{R}$ respectively as the average variance across traits of shared and trait-specific effects across traits.

##### 2.2.3 Phenotype Predictions

The LIMIX variance decomposition model also implements a tool for phenotype predictions, which can be used to monitor overfitting and perform model selection. For example, this can help select the set of fixed and random effects to consider, the best parametrisation of the trait-to-trait covariance matrices, or the best extent of the matrix penalization.

Following (1.14), the predictive posterior mean of the contribution all terms in model (2.7) is

$$\text{vec}(\boldsymbol{M}^\star) = \sum_i \left(\boldsymbol{A}_i \otimes \boldsymbol{X}_i^\star\right) \text{vec}(\boldsymbol{B}_i) + \sum_i \left(\boldsymbol{C}_i \otimes \boldsymbol{R}_i^\star\right) \boldsymbol{K}^{-1} \text{vec}\left(\boldsymbol{Y} - \sum_i \left(\boldsymbol{A}_i \otimes \boldsymbol{X}_i\right) \text{vec}(\boldsymbol{B}_i)\right) \tag{2.8}$$

where $\boldsymbol{K} = \sum_i \boldsymbol{C}_i \otimes \boldsymbol{R}_i + \boldsymbol{\Sigma} \otimes \boldsymbol{I}$ and $\boldsymbol{R}_i^\star$ is the cross covariance for term $i$. For example, if $\boldsymbol{R}$ has the form in (2.6) then we have $\boldsymbol{R}^\star \propto \boldsymbol{W}_{:,\mathcal{S}} {\boldsymbol{W}^\star_{:,\mathcal{S}}}^T \in \mathbb{R}^{NN^\star}$ where N and $N^\star$ are the number of samples in the training and test sets respectively and $\boldsymbol{W}$ and $\boldsymbol{W}^\star$ are their genotype.

#### 2.3 Genome-wide Association Studies

In this section we briefly describe the methods implemented in LIMIX to perform single and multi-locus analysis in GWAS.

##### 2.3.1 univariate GWAS

The standard LMM considered in GWAS is

$$\boldsymbol{y} \sim \mathcal{N}\left(\boldsymbol{W}\boldsymbol{\alpha} + \boldsymbol{x}\beta,\, \sigma_g^2\left(\boldsymbol{R} + \delta\boldsymbol{I}\right)\right) \tag{2.9}$$

where $\boldsymbol{x}$ is the genotypic profile of the SNP being tested, $\beta$ its effect size, $\boldsymbol{R}$ the sample-to-sample relatedness matrix and $\delta = \sigma_g^2/\sigma_e^2$ the signal-to-noise ratio. Testing for association of the SNP to the phenotype is done by testing $\beta \neq 0$. The $\delta$ representation of the marginal likelihood allows fast scalable methods and has been widely used in GWAS [Lippert et al., 2011, Zhou and Stephens, 2012].

P: not sure we want to say this The variance decomposition tool from LIMIX can be used to build more complex forms of the relatedness matrix $\boldsymbol{R}$. For example, considering the model (2.5) introduced in the previous section, we can estimate the variance components $\{\hat{\sigma}_i\}_i$ by maximum likelihood or penalized maximum likelihood and then define a new relatedness matrix $\boldsymbol{R} = \sum_i \hat{\sigma}_i^2 \boldsymbol{R}_i / \sum_i \hat{\sigma}_i^2$ that accounts for all contributions. This is equivalent to fix the contributions of each random effect term on the null model while the signal-to-noise ratio is still flexibly evaluated SNP-wise using the model in (2.9). However, building a too complex background model might explain away genetic signal leading to power reduction in GWAS. In all our experiments we just considered the full kinship matrix as relatedness matrix, with the exception of the transcript-based eQTL analysis where we accounted for hidden confounders by using PANAMA (section 2.4).

##### 2.3.2 multivariate GWAS

The multivariate version of 2.9 is

$$\boldsymbol{Y} \sim \mathcal{N}\left(\left(\boldsymbol{A}_{\text{cov}} \otimes \boldsymbol{W}\right)\text{vec}\left(\boldsymbol{B_0}\right) + \left(\boldsymbol{A}_1 \otimes \boldsymbol{x}\right)\text{vec}\left(\boldsymbol{B}\right),\, \sigma_g^2\left(\boldsymbol{C} \otimes \boldsymbol{R} + \delta\boldsymbol{\Sigma} \otimes \boldsymbol{I}\right)\right) \tag{2.10}$$

where the trait-trait matrices $\boldsymbol{C}$ and $\boldsymbol{\Sigma}$ can be estimated using the variance decomposition tool and then plugged in. This equals to estimate the variance parameters on the model with no association with the marker. However, LIMIX allows for SNP-specific estimates of the global variance of random effects $\sigma_g^2$ and the signal-to-noise ratio $\delta$. The model in (2.10) is used to test for specific trait designs of the marker on the multivariate phenotype. To do so, the alternative model in (2.10) is compared to a null model which has the same form but characterized by a different trait design $\boldsymbol{A}_0$ for the marker effect. In the following we describe the choices of $\boldsymbol{A}_0$ and $\boldsymbol{A}_1$ to conduct the tests that are most commonly considered in multivariate GWAS and some of their extentions.

- *Any effect test*: this is a $P$ degrees of freedom test where we test if the marker has an effect on at least one of the phenotypes ($\boldsymbol{A}_1 = \boldsymbol{I}_P$ and $\boldsymbol{A}_0 = \boldsymbol{0}_P$).
- *Common effect test*: this is a one degree of freedom test, where in the alternative model the marker has the same effect size and direction across all phenotypes ($\boldsymbol{A}_1 = \boldsymbol{1}_P$) while the null model does not contain the effect from the marker ($\boldsymbol{A}_0 = \boldsymbol{0}_P$).
- *Specific effect test*: this is a one degree of freeform test that we can use to test if the genetic marker acts specifically on the phenotype $p$. The design in the alternative model is a combination of a common effect and an independent effect for trait $p$ $\boldsymbol{A}_1 = [\boldsymbol{1}_P, \boldsymbol{1}_{\text{trait}==p}]$, where $\boldsymbol{1}_{\text{trait}==p}$ return a vector with 1 at element $p$ and 0 at all other elements, while in the null model we just have the common effect, $\boldsymbol{A}_0 = \boldsymbol{1}_P$.
- *Any specific effect test*: this is a $P - 1$ degrees of freeform test where we test whether the marker has a specific effect on at least one of the phenotypes. The alternative model describes an *any effect* from the marker ($\boldsymbol{A}_1 = \boldsymbol{I}_P$) while the null model describes a *common effect* ($\boldsymbol{A}_0 = \boldsymbol{1}_P$).

LIMIX flexibly allows considering more complex tests to really exploit the information in multivariate datasets. Here we describe an example of a more complex test. Suppose we want to jointly model $T$ traits across $E$ different environments, for a total of $P = TE$ trait-environment combinations. In this example, we might be interested in discovering loci that affect multiple phenotypes differently across different environment. This is a $T$ degree of freedom test and can be performed in LIMIX by considering an alternative model where the marker has different effect sizes across traits and environments ($\boldsymbol{A}_1 = \boldsymbol{I}_P$) and a null model where the marker has same effect size across different environments for each trait ($\boldsymbol{A}_0 = \boldsymbol{I}_T \otimes \mathbf{1}_{1,E}$).

P: not sure we want to leave this in LIMIX makes also available a more general model for multivariate GWAS that handles non-Kronecker structured fixed effects and incomplete designs. However, such a model does not allow speed-ups and can be only used for the joint analysis of a few traits. The model implemented for such a task is practically the analogous of (2.9):

$$\mathrm{vec}\left(\boldsymbol{y}\right) \sim \mathcal{N}\left(\boldsymbol{W}\boldsymbol{\alpha} + \boldsymbol{a}_0 \odot \boldsymbol{x}\beta_0 + \boldsymbol{a}_1 \odot \boldsymbol{x}\beta_1,\ \sigma_g^2\left(\boldsymbol{K} + \delta\boldsymbol{I}\right)\right) \tag{2.11}$$

where $\boldsymbol{y}$ is an $NP \times 1$ multi-phenotype vector, $\boldsymbol{y} = \mathrm{vec}\left(\boldsymbol{Y}\right)$, and an element-wise product $\odot$ is introduced to introduce flexible designs. While the first fixed effect for SNP $\boldsymbol{x}$ is contained both in the full and the null model, we test for design $\boldsymbol{a}_1$ by testing $\beta_1 \neq 0$. For example, if $\boldsymbol{l}$ labels a categorical variable with values in $\{l_1, l_2, l_3\}$, setting $\boldsymbol{a}_0 = \mathbf{1}_P$ and $\boldsymbol{a}_1 = \mathbf{1}_{\boldsymbol{l}==l_1}$ we can test for $l_1 \times$SNP interactions.

##### 2.3.3 Multi-locus GWAS

All the models we introduced in this section can be extended to consider multiple loci by step-wise forward selection [Segura et al., 2012]. This is the first time multivariate analysis and step-wise forward selection are combined together. The general framework LIMIX employs to perform multi-locus GWAS encompasses two main steps

1. a genome-wide scan is performed using the model in (2.10)
2. if the most associated marker has P-value lower than a certain threshold the marker is added as covariate and back to step 1, otherwise the algorithm stops here.

LIMIX allows the user to

- set the type of test to be performed in the genome-wide scans;
- set the design of the SNPs that are added as covariates;
- set a threshold either over the P-value or the Q-value of the most associated SNP as stopping criterion;
- add a maximum number of iterations as stopping criterion;
- update the parameters of the trait-to-trait covariance matrices after each inclusion using internally the variance decomposition module.

#### 2.4 PANAMA

The PANAMA tool from LIMIX can be used to find a sample-by-sample covariance matrix that accounts for genetic and non-genetic confounders [Fusi et al., 2012]. This can be done when a high number of traits are thought to share common confounding structure. This is the case of eQTL analysis, where thousands of genes most likely share sample-specific confouders. Reproducing the framework used in gaussian process latent variable model [Lawrence, 2004, 2005], we assume independence of the traits (e.g., gene expression in eQTL analysis) conditions on the latent factors. After normalizing all traits to z-scores, we again consider a particular model from the general form (2.1):

$$\boldsymbol{y}_p = \boldsymbol{u} + \boldsymbol{\eta} + \boldsymbol{\psi}, \quad \forall p \tag{2.12}$$

$$\boldsymbol{u} \sim \mathcal{N}\left(\mathbf{0},\ \sigma_g^2 \boldsymbol{R}_{\mathrm{geno}}\right),\ \boldsymbol{u} \sim \mathcal{N}\left(\mathbf{0},\ \boldsymbol{R}_{\mathrm{conf}}\left(\boldsymbol{\alpha}\right)\right),\ \boldsymbol{\Psi} \sim \mathcal{N}\left(\mathbf{0},\ \sigma_e^2 \boldsymbol{I}\right)$$

where $\boldsymbol{R}_g$ is the genetic relatedness matrix introduced before while $\boldsymbol{R}_c(\boldsymbol{\alpha})$ is the relatedness matrix due to unobserved covariates. The parameters of the model are $\{\boldsymbol{\alpha}, \sigma_g^2, \sigma_e^2\}$. As full rank matrices might overfit the data explaining away genetic signal, LIMIX models $\boldsymbol{R}_c(\boldsymbol{\alpha})$ as rank $r$ matrix and allows the user to choose $r$. A sensible value for $r$ can be chosen by looking at the number of PCAs of the sample-by-sample empirical covariance matrix explaining 70%-90% of the variance.

Once $\boldsymbol{R}_c$ is learned from the data, a new sample-to-sample relatedness matrix can be plugged in the GWAS tools both for single and multivariate analysis.
