## Supplementary material for "LIMIX: genetic analysis of multiple traits": Data Supplement 1 (from another version of this preprint: version 1, 2014-05-20)

### Supplemental Information
### Multivariate analysis of heritable traits

Christoph Lippert*,† Franceso Paolo Casale*, Barbara Rakitsch, Oliver Stegle*,†

#### 1 NFBC

**Supplementary Table 1 : Tabular summary of significant associations in the NFBC datasets identified using alternative LMM methods.** This file is included as separate supplementary file (supplementary_Table1.xls).

*These authors have contributed equally.

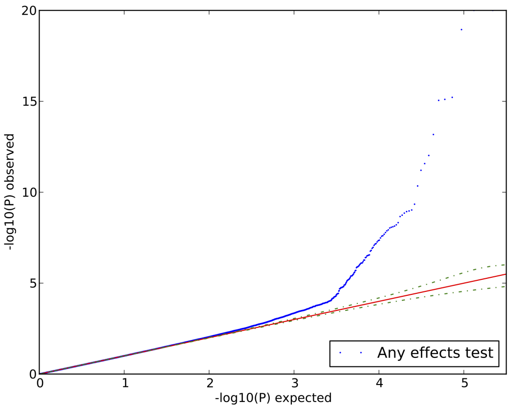


**(a)** Any effect test

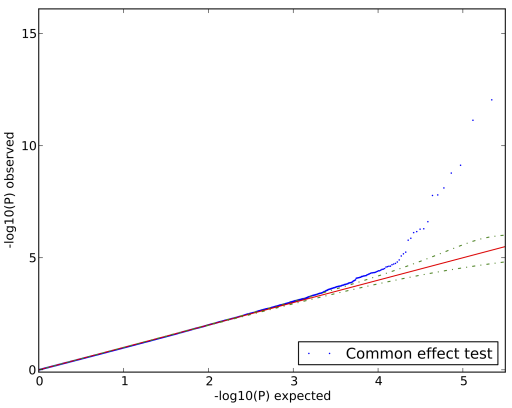


**(b)** Common effect test

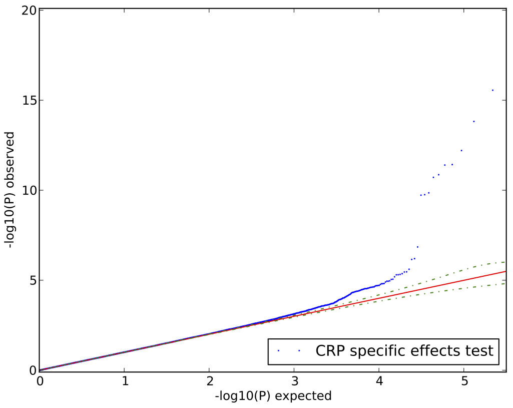


**(c)** CRP specific effect test

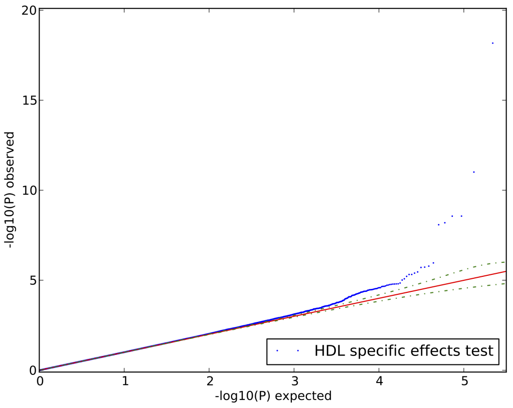


**(d)** HDL specific effect test

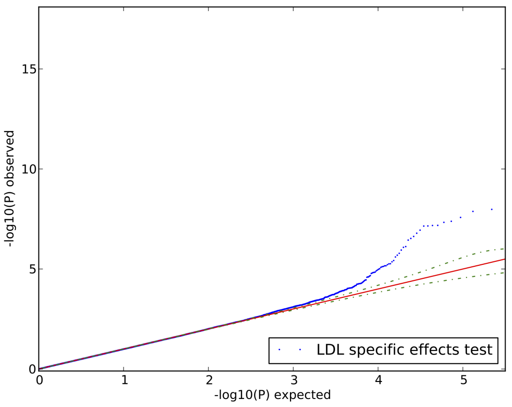


**(e)** LDL specific effect test

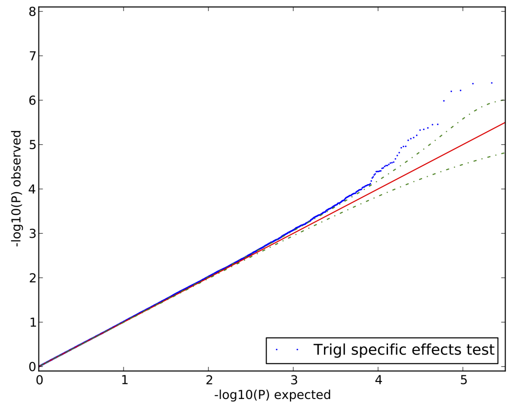


**(f)** Trigl specific effect test

**Supplementary Figure 1 $Q-Q$-plots for the multi-trait association tests on the NFBC dataset.** Shown are $Q-Q$-plots for the any effect test, common effect test and specific effect test for each of the four phenotypes (CFP,HDL,LDL and TRIGL).

### 2 Geuvadis

**Supplementary Table 2 : Tabular summary of *cis* associations identified in the Geuvadis dataset considering alternative multi-trait LMMs and tests.** This file is included as separate supplementary file (supplementary_Table2.xlsx).

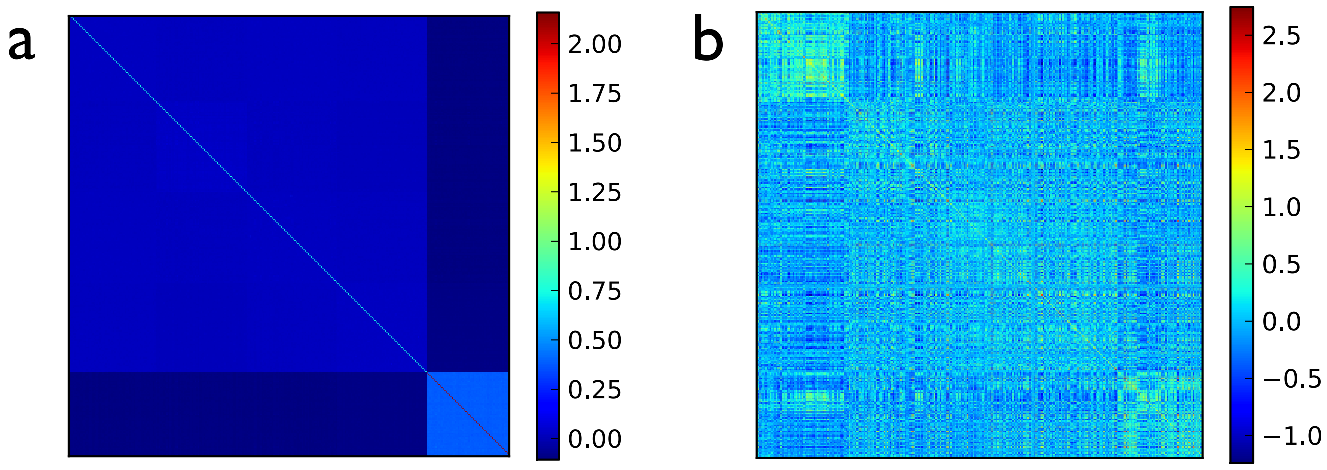


**Supplementary Figure 2 Sample covariance matrices used in mixed model analyses, either estimated from genetic kinship or using the PANAMA model in the human eQTL dataset.** Shown is the genetic Kinship covariance estimated using a linear kernel (**a**) and an equivalent matrix estimated using the PANAMA model (**b**). The panama model uses both genotype data and expression levels for the covariance estimate, thereby accounting for hidden confounding factors. Samples are grouped by population (CEU, FIN, GBR, TSI, YRI).

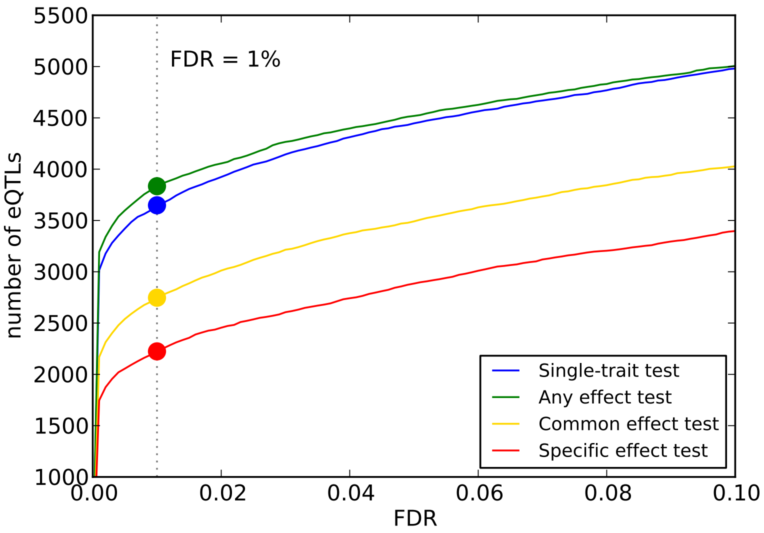


**Supplementary Figure 3 Power comparison among different trait design tests on the human eQTL dataset.** Compared were the any effect test, common effect test and specific effect test for individual isoforms as well as marginal analysis. Shown are the number of genes with at least one significant eQTL of a particular type and for increasing false discovery rate cutoffs (x-axis).

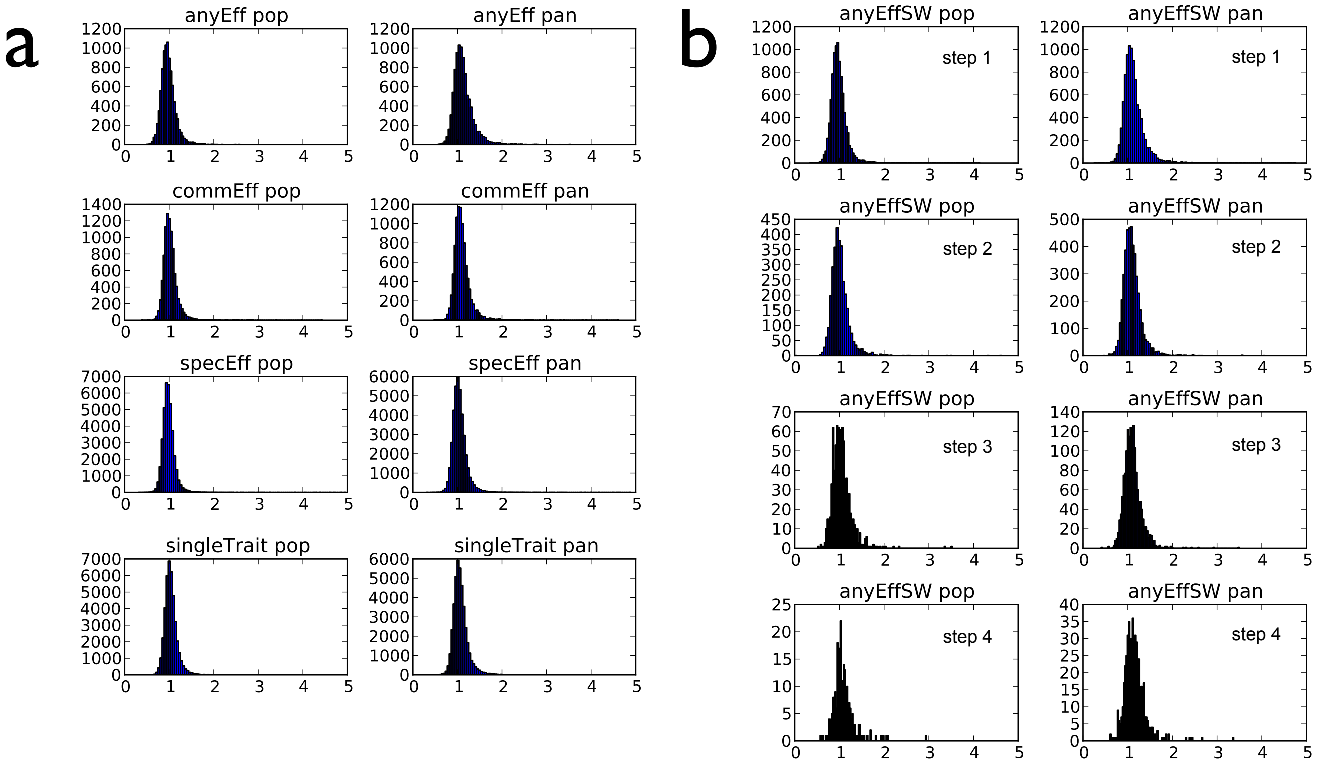


**Supplementary Figure 4 Distribution of genomic control for the *cis* eQTL association analyses in the human eQTL dataset.** Histogram show $\lambda_{GC}$ estimates obtained for tests considered in the human eQTL analysis, either using a sample covariance inferred from genotype kinship (pop) or using the PANAMA model (pan). **a,** histograms for any effect test, common effect test, specific effect tests and single trait association test (*anyEff*, *commEff*,*specEff*,*singleTrait* respectively) while **b** shows histograms for steps 1-4 of multi trait multi locus analysis (*anyEffSW*). The results show that both the PANAMA and the standard kinship-based sample covariance yield calibrated test statistics, as do the multi-locus association analyses carried out.

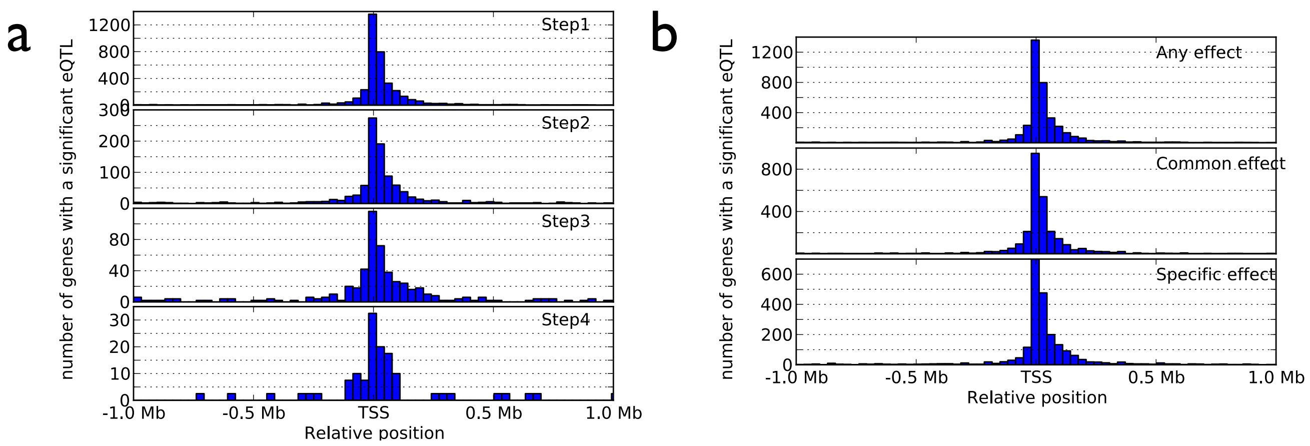


**Supplementary Figure 5 Distribution of the relative position of eQTLs identified using alternative multi-trait LMMs in the human eQTL dataset.** The histograms show the position of the most associated variant relative to the transcription start site of the corresponding gene for significant eQTLs (FDR $\leq$ 1%), considering alternative methods and tests. **a,** different association steps in the multi-locus multi-trait model when considering the any effect test. Shown are histograms for the primary association (step 1), secondary association (step 2) or higher-order association signals (step 3,4). **b,** primary associations of the multi-trait model, considering either the any effect test (as in **a**, step 1), gene-level effects (common effect test) or genetic effects that are specific to individual isoforms (specific effect test). In sum, eQTLs identified by all methods tended to occur in the vicinity of the transcription start site, suggesting the associations are genuine. Specific effect tests and higher-order association signals (step 4) tended to be slightly more likely to occur in distal regions.

#### 3 Yeast

**Supplementary Table 3 : Tabular summary of the average variance contribution in yeast pathways estimated using the variance decomposition model.** This file is included as separate supplementary file (supplementary_Table3.xlsx).

**Supplementary Table 4 : Tabular summary of significant associations on the *lysine bios.* yeast pathway.** This file is included as separate supplementary file (supplementary_Table4.xlsx).

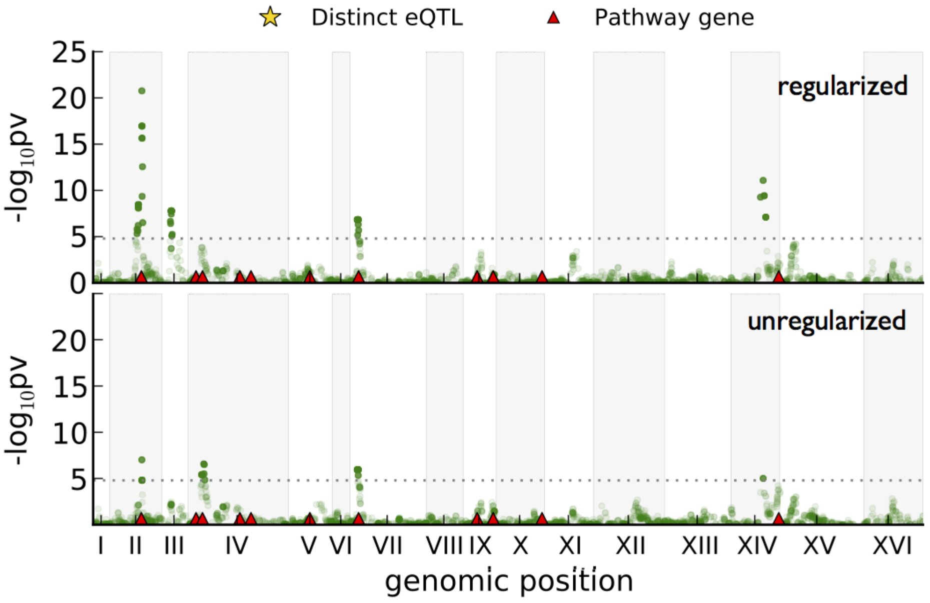


**Supplementary Figure 6 Power comparison of regularized and unregularized multi-trait LMMs for the analysis of the *lysine bios.* pathway in the yeast eQTL dataset.** Shown are Manhattan plots for the optimally regularized LMM (top panel) and a standard unregularized LMM (lower panel) using an any effect association test applied to the 22 traits (11 genes, 2 environments) of the *lysine bios.* pathway in yeast.

| Step | Pval MT | Pval GxE | Pval Common | SNP pos | SNP chrom | closest gene in the p | type | closest gene | gene TSS | dist (Kb) | Pval ST |
|---|---|---|---|---|---|---|---|---|---|---|---|
| 1 | 1.72E-21 | 4.90E-03 | **1.39E-21** | 477206 | 2 | YBR115C | CIS | YBR117C | 476431 | 0.775 | 2.47E-21 |
| 2 | 6.10E-14 | 4.69E-04 | **3.38E-12** | 486861 | 14 | YNR050C | TRANS | YNL076W | 483557 | 3.304 | 9.87E-11 |
| 3 | 5.51E-12 | 2.87E-02 | **2.39E-12** | 92013 | 3 | - | TRANS | YCR027C | 167995 | 75.982 | 7.27E-06 |
| 4 | 2.10E-11 | 9.40E-01 | **6.02E-15** | 98231 | 7 | YGL208W | CIS | YGL208W | 97342 | 0.889 | 7.93E-10 |
| 5 | 5.04E-12 | **2.18E-07** | **1.38E-06** | 174364 | 15 | - | TRANS | YOL077W-A | 185438 | 11.074 | - |
| 6 | 6.08E-08 | 4.66E-03 | **7.62E-07** | 217351 | 4 | YDL131W | CIS | YDL137W | 216529 | 0.822 | 1.34E-05 |
| 7 | 1.13E-06 | 3.40E-03 | 2.76E-05 | 579459 | 2 | YBR115C | TRANS | YBR170C | 578081 | 1.378 | - |
| 8 | 4.22E-07 | **1.03E-08** | 4.04E-01 | 446685 | 11 | - | TRANS | YKR003W | 445024 | 1.661 | - |
| 9 | 6.18E-06 | 1.10E-01 | **2.34E-06** | 242417 | 9 | YIL094C | CIS | YIL064W | 241940 | 0.477 | - |
| 10 | 1.48E-07 | 3.53E-01 | **4.02E-09** | 403134 | 9 | YIR034C | CIS | YIR024C | 403488 | 0.354 | - |

**Supplementary Table 5 : Tabular summary of significant associations identified in the *lysine bios.* pathway in the yeast dataset using alternative LMM methods.** This file is in addition provided as separate supplementary .xls file (SupplementaryResultsYeast.xls).

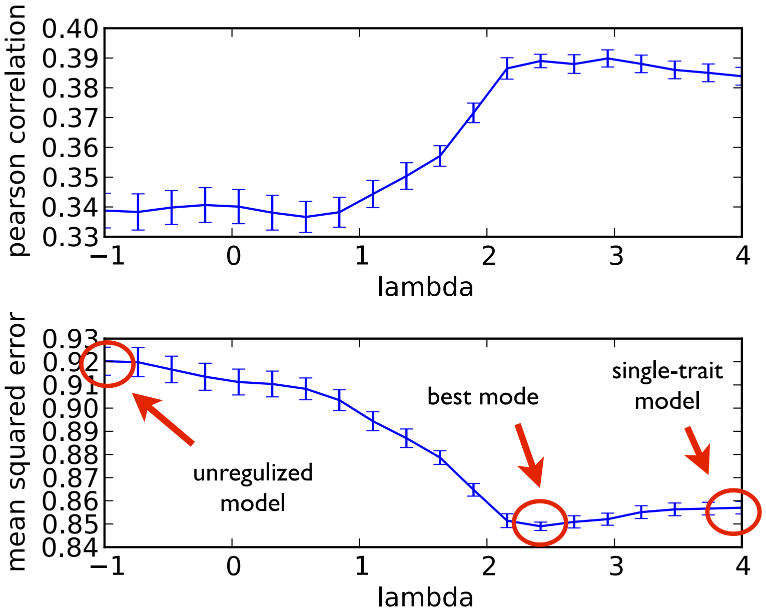


**Supplementary Figure 7 Model selection approach to determine optimal regularization of the multi-trait LMM applied to the *lysine bios.* pathway in the yeast eQTL dataset.** Model selection is carried out using out-of-sample prediction (10-fold cross validation). Shown are out-of-sample correlation coefficients (top panel) and mean-squared errors (lower panel) for the corresponding selection experiments. The ideal model is determined by minimizing the prediction error, which is lower than those obtained from a fully unregularized multi-trait model (far left) or a single-trait model (far right).

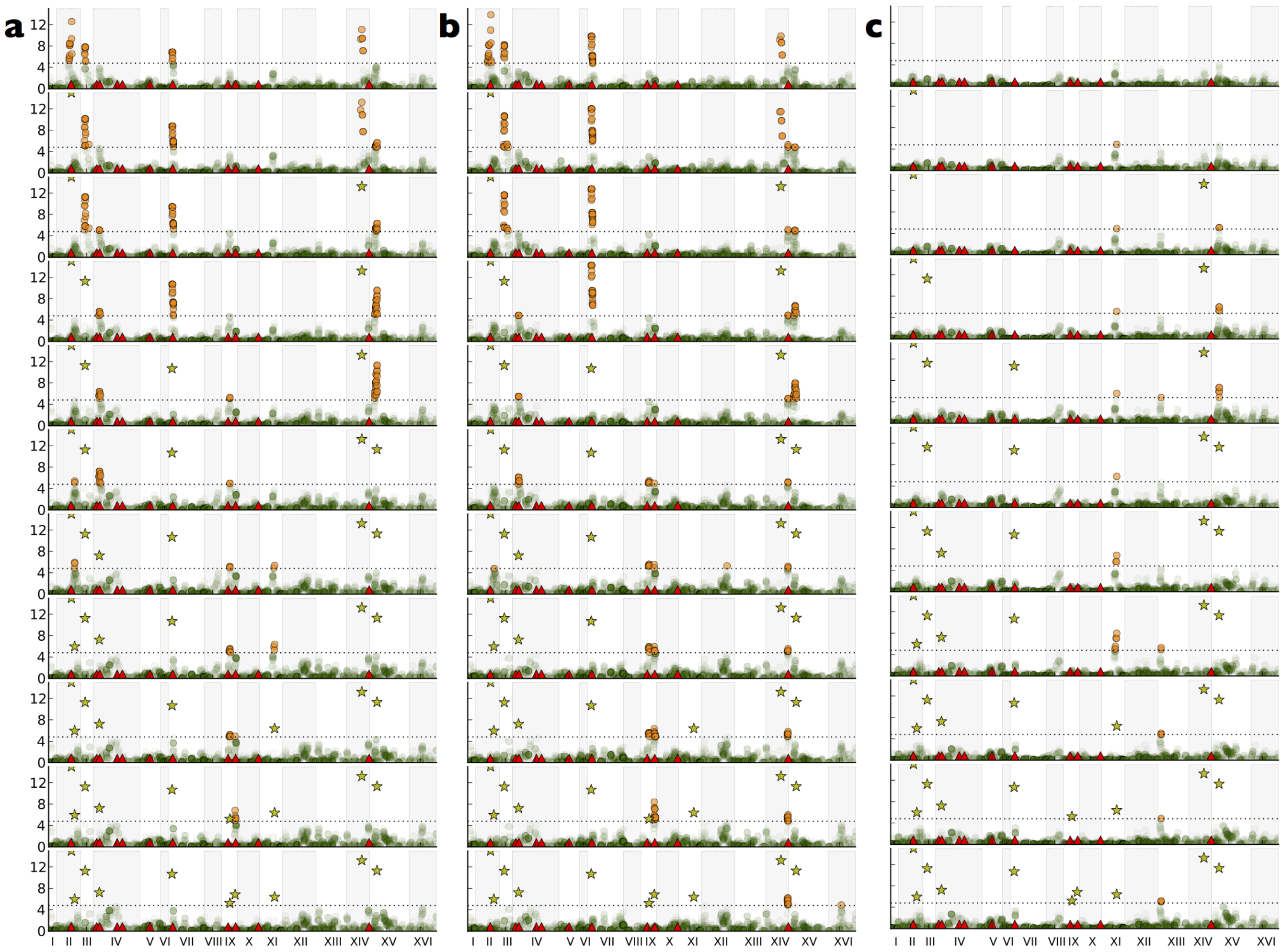


**Supplementary Figure 8 Illustration of the step-wise inclusion of fixed effects used for multi-trait multi-locus LMMs applied to the *lysine bios.* pathway in the yeast eQTL dataset.** Considered was the multi-trait multi-locus model at different iterations of the stepwise inclusion of fixed effect SNPs (from top to bottom). Alternative tests using the multi-trait LMM were considered, **a,** any effect test, **b,** common effect test and **c,** GxE effect test. Stepwise inclusion of individual loci was carried out using the any effect test, whereas the alternative tests were used to annotated loci included in the model. The GxE analysis corresponds to a specific effect-test for the ethanol condition (one of two environments), where this 11 degrees of freedom test couples the 11 genes. Briefly, while the majority of identified eQTLs were significant under the common effect test (suggesting persistence across environments), there were two significant GxE effects on chromosomes *XI* and *XV*.

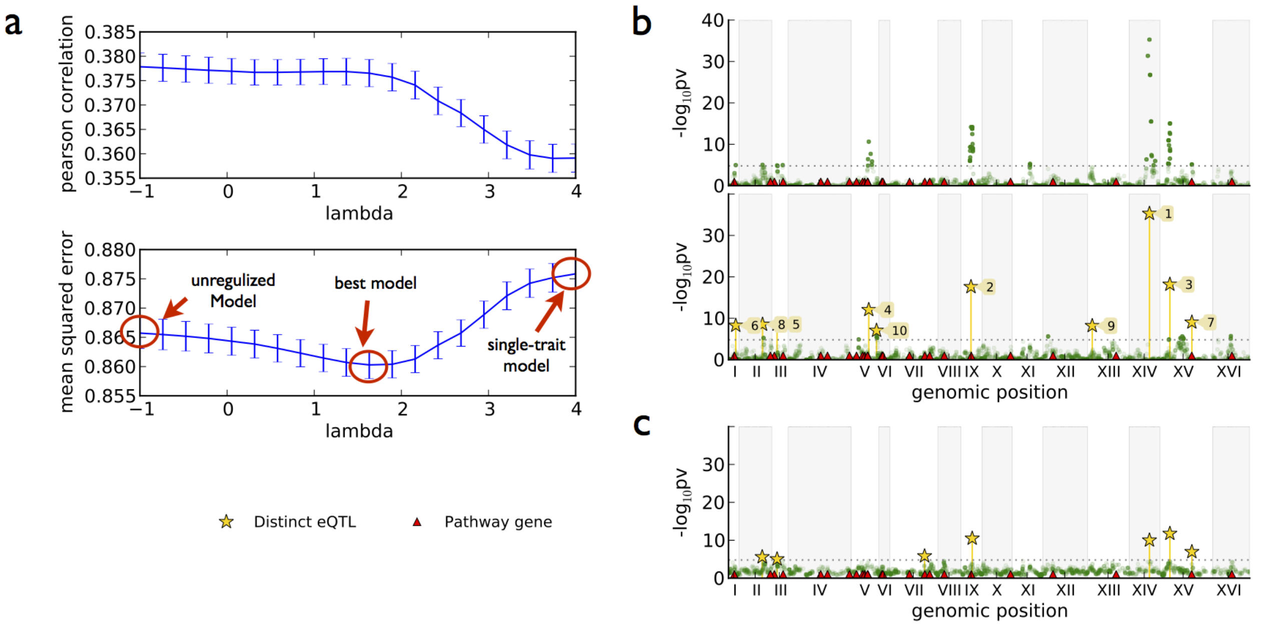


**Supplementary Figure 9 Multi-trait LMM analyses of 24 genes in the *glycine, s. and t. metabolism* pathway across two environments (48 traits) in the yeast eQTL dataset. a,** model selection using model out-of-sample prediction, yielding an optimally regularized model for genetic analysis (see also Supplementary Fig. 7). **b,** multi-locus analysis, showing the first 10 significant associations identified by the optimally regularized multi-trait LMM. Upper panel: step 1, corresponding to a multirait-LMM; lower panel: step 10. **c,** equivalent results using a single-trait multi-locus model, where only 7 significant associations are found. In sum, these results confirm the benefits of the multi-trait multi-locus analysis on a second independent pathway.

### 4 Validation and prediction

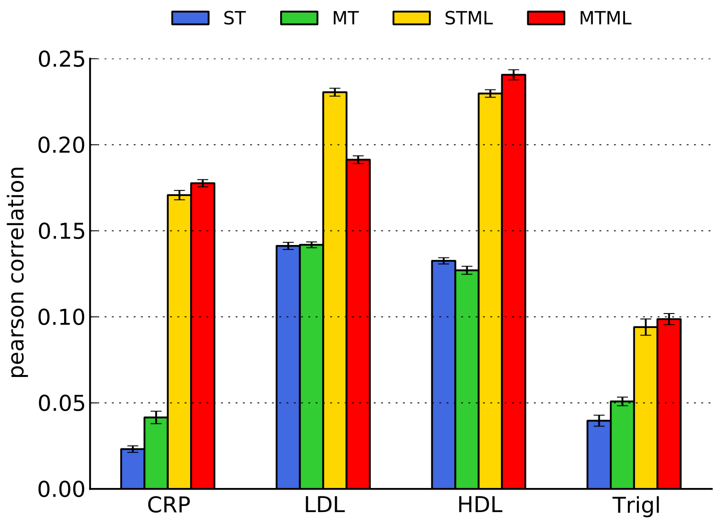


**Supplementary Figure 10 Assessment of alternative LMMs for out-of-sample prediction on the NFBC dataset.** Model comparison using out-of-sample prediction (ten repeats of 5-fold cross validation). Compared were a BLUP-based single-trait model (ST), a multi-trait equivalent model (MT) and multi-locus equivalents (STML and MTML), where in addition to the random effect term, individual SNPs have been iteratively selected as fixed effects (Methods). All model parameters were fit on the training data alone, including the selection of fixed effect SNPs for the multi-locus models. Multi-locus LMMs (single trait or multi trait) performed consistently better than the corresponding LMMs without fixed effects (STML vs ST & MTML vs MT). On the whole, multivariate models were found to make more accurate predictions than univariate approaches, where the combination of multi-locus and multi-trait LMM achieved best prediction performance overall.

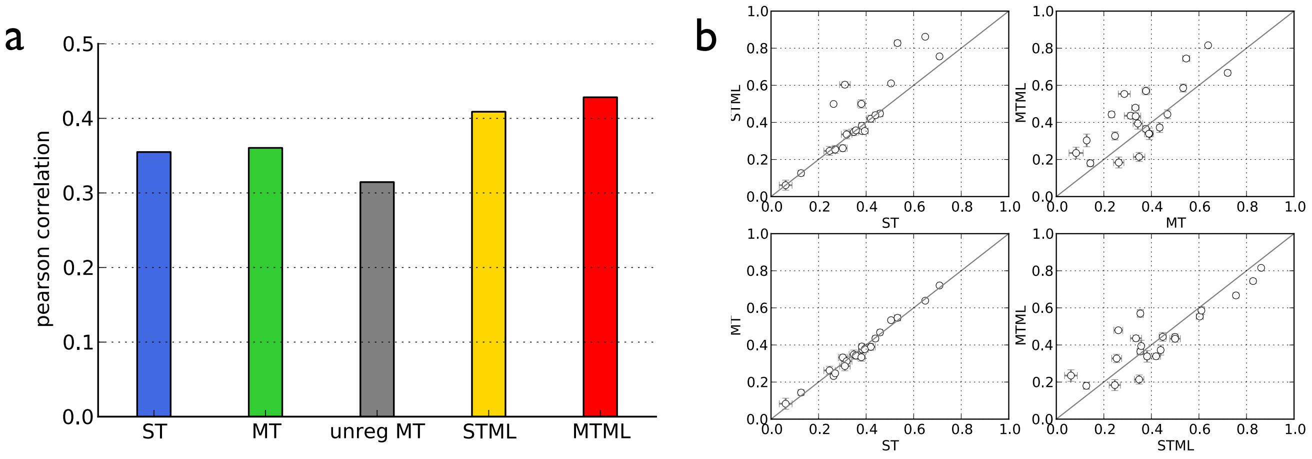


**Supplementary Figure 11 Assessment of alternative LMMs for out-of-sample prediction of genes in the *lysine bios.* pathway in the yeast eQTL dataset.** Compared were a BLUP-based single-trait model (ST), a multi-trait equivalent model (MT) and multi-locus equivalents (STML and MTML), where in addition to the random effect term, individual SNPs have been iteratively selected as fixed effects (Methods). All model parameters were fit on the training data alone, including the model-selection for optimal regularization (see Supplementary Figure 7) and the selection of fixed effect SNPs for the multi-locus models. For comparison, an unregularized mixed model was also considered (unreg MT), where the model selection step was omitted. **a,** average out-of-sample Pearson correlation coefficient for 10 repeat experiments of 10-fold cross validation, using the considered models for prediction. **b,** scatter plot of Pearson correlation coefficients for individual genes (between predicted and real gene expression levels), comparing alternative methods pairwise. The results show that multi-trait modeling is superior to single-trait models (MT vs ST) and that multi-locus models yield improved prediction accuracy (STML versus ST and MTML versus MT). Unregularized multi-trait models (unreg MT) perform worse than single-trait models, demonstrating the need for appropriate model selection.

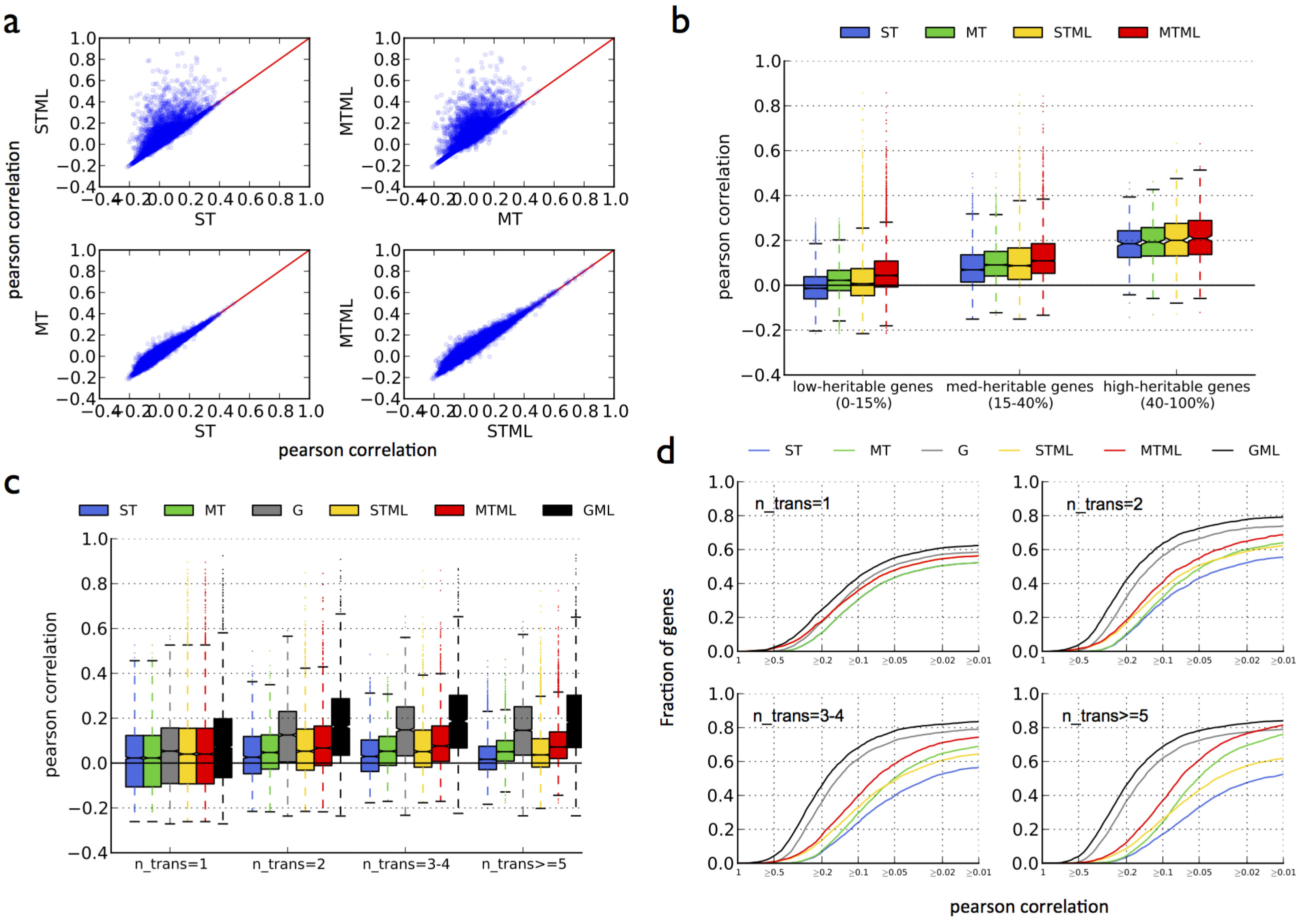


**Supplementary Figure 12 Assessment of alternative LMMs for out-of-sample prediction of isoform and gene expression levels in the human eQTL dataset.** Prediction experiments were carried out using 10-fold cross validation for out-of-sample prediction. For each gene, the average Pearson correlation coefficient (between predicted and measured isoform expressions) across isoforms of each gene is reported. All genes expressing 2-10 transcript isoforms (9,246 total) were considered for analysis. Compared were a BLUP-based single-trait model (ST), a multi-trait linear mixed model model (MT) and multi-locus equivalents (STML and MTML), where in addition to the random effect term, individual SNPs have been iteratively selected (Online Methods). For comparison, we also considered classical gene expression estimates as phenotype for prediction (in **c**). Here, the full set of 15,220 genes with at least one expressed isoform was used for analysis (Online Methods). **a,** prediction accuracy of different methods, comparing genome-wide out-of-sample prediction accuracy; **b,** identical results, stratified by estimated heritability of individual genes. **c,** prediction accuracy stratified by the number of expressed isoforms for each gene and **d,** showing cumulative fractions of genes with decreasing prediction accuracies. The differences in prediction performance for gene-level expression estimates and transcript-level prediction even for genes with a single expressed transcript (**c,d**) arise from the filtering procedure applied to remove unexpressed transcripts (Online Methods).
